# Patient-derived extracellular matrix demonstrates role of COL3A1 in blood vessel mechanics

**DOI:** 10.1101/2022.10.16.512399

**Authors:** Elizabeth L. Doherty, Wen Yih Aw, Emily C. Warren, Max Hockenberry, Grace Krohn, Stefanie Howell, Brian O. Diekman, Wesley R. Legant, Hadi Tavakoli Nia, Anthony J. Hickey, William J. Polacheck

**Affiliations:** Joint Department of Biomedical Engineering, University of North Carolina at Chapel Hill and North Carolina State University; UNC Catalyst for Rare Diseases, Eshelman School of Pharmacy, University of North Carolina at Chapel Hill; Department of Cell Biology and Physiology, University of North Carolina at Chapel Hill School of Medicine; Department of Pharmacology, University of North Carolina at Chapel Hill School of Medicine; Department of Biomedical Engineering, Boston University; McAllister Heart Institute, University of North Carolina at Chapel Hill

**Keywords:** Vascular Ehlers-Danlos syndrome, cell derived matrix, collagen III, mechanotransduction, ECM mechanics, viscoelasticity

## Abstract

Vascular Ehlers-Danlos Syndrome (vEDS) is a rare autosomal dominant disease caused by mutations in the *COL3A1* gene, which renders patients susceptible to aneurysm and arterial dissection and rupture. To determine the role of *COL3A1* variants in the biochemical and biophysical properties of human arterial ECM, we developed a method for synthesizing ECM directly from vEDS donor fibroblasts. We found that the protein content of the ECM generated from vEDS donor fibroblasts differed significantly from ECM from healthy donors, including upregulation of collagen subtypes and other proteins related to ECM structural integrity. We further found that ECM generated from a donor with a glycine substitution mutation was characterized by increased glycosaminoglycan content and unique viscoelastic mechanical properties, including increased time constant for stress relaxation, resulting in a decrease in migratory speed of human aortic endothelial cells when seeded on the ECM. Collectively, these results demonstrate that causal *COL3A1* mutations lead to the synthesis of ECM that differs in composition, structure, and mechanical properties from healthy donors. These results further suggest that ECM mechanical properties could serve as a prognostic indicator for patients with vEDS, and the insights provided by the approach demonstrate the broader utility of cell-derived ECM in disease modeling.

## 1. Introduction

Vascular Ehlers-Danlos syndrome (vEDS, Ehlers-Danlos Syndrome Type IV) is a rare autosomal dominant disorder caused by mutations in the *COL3A1* gene [1]. There are no US Food and Drug Administration (FDA) approved therapies for the treatment of vEDS and the mortality rate after surgical intervention remains high [2, 3]. Unpredictable spontaneous rupture of hollow organs, including the arterial vasculature, uterus, and intestines can occur at a young age [4], and arterial rupture is the leading cause of death [3]. In addition to the formation and rupture of aneurysms, which are among the most common complications, patients are at risk of spontaneous rupture of nonaneurysmatic vessels [2]. Further, the risk of vessel rupture is widely distributed throughout arterial vasculature [5], demonstrating systemic arterial fragility. Recent data from clinical trials have demonstrated that administration of β-adrenergic agonist blockers mitigates risk for arterial rupture by reducing systemic blood pressure and hemodynamic stress at the arterial wall [6, 7] and suggests that addressing the pathologic mechanics of afflicted arteries could be a successful strategy for treatment. However, the mechanisms by which the causal *COL3A1* variants contribute to increased risk of vascular rupture remain unclear.

Previous work has sought to establish a connection between *COL3A1* mutations and the mechanical properties of arterial tissue in vEDS patients and in murine models exhibiting *COL3A1* mutations. *In situ* measurements using ultrasound and blood pressure monitoring did not detect differences in the mechanical properties of the carotid artery for patients with vEDS [8, 9]. Furthermore, while a significant reduction of the intima-media thickness and elevated stress in the arterial wall was observed in vEDS patients [8], subsequent investigation using magnetic resonance imaging (MRI) to image carotid artery wall thickness including the adventitia did not identify any differences in arterial wall thickness in patients with vEDS [10]. Pulse wave analysis suggests a slight increase in arterial wall stiffness, though these studies evaluated multiple EDS subtypes [11, 12]. Murine models have demonstrated a decrease in the tensile force at rupture of murine thoracic aortic explants [13], but data from murine models is difficult to interpret in the context of vEDS pathophysiology because murine models fail to recapitulate key aspects of the disease. For example, mice that are haploinsufficient for *COL3A1* do not display a clinically observable vascular phenotype or differences in morbidity and mortality as compared to wild-type control mice [14, 15], while the human phenotype exhibits a severe pathology, leading to significant morbidity due to vascular rupture [16]. Furthermore, the regulation of gene networks of *COL3A1* differ significantly between mice and humans [17], which presents a significant challenge for translating drug targets and therapies identified in mice to the clinic. Collectively, these studies fail to demonstrate a clear connection between causal *COL3A1* variants and biomechanical phenotypes of arterial tissue of patients with vEDS.

Collagen III, which is formed from a triple helix comprised of three α1(III) chains encoded by the *COL3A1* gene [18], is a highly abundant fibrillar collagen enriched in hollow organs, including blood vessels, with collagen I and collagen III being the most abundant collagens in the vascular wall [19]. While collagen III forms fibrils independent of collagen I [20], these fibrils are relatively compliant compared to collagen I fibrils [21], and the effects of collagen III on ECM and tissue mechanics are largely driven by its role as a regulatory protein in the fibrillogenesis of collagen I [22]. *In vitro*, the presence of collagen III reduces collagen I fibril diameter and resulting elastic modulus of composite fibrils in a dose-dependent manner [21]. Consistent with these results, heterozygous *COL3A1* knockout mice (*COL3A1*^*+/-*^) demonstrate increased collagen I fibril diameter in cartilage tissue, though in contrast to the fibril-level mechanics, the elastic modulus of cartilage tissue is reduced with reduction of collagen III [23]. A possible explanation for the difference in fibril-level vs. tissue-level mechanics is the difference in fibril crosslinking observed in *COL3A1*^*+/-*^ mice [23]. Furthermore, transcriptome analysis of vEDS patient-derived cells have demonstrated altered expression levels of ECM-related genes including *MMP24* (upregulated), and *FNB2* and *LOXL3* (downregulated) [24], and previous studies also show that total ECM secretion by vEDS patients is reduced compared to cells from healthy patients [24-26]. Thus, it remains unclear whether defective collagen III or subsequent expression of other matrix-associated genes drives the clinical phenotype, as in certain populations with classical EDS [27].

Here, we investigated the contribution of *COL3A1* mutations to cell and ECM mechanics intending to further characterize the role of *COL3A1* in arterial fragility and aneurysm risk in patients with vEDS. To enable these studies and to address the limitations of murine vEDS models, we developed a technique for generating collagen-rich ECM from patient donor fibroblasts. To account for variability in vEDS causal mutations, we generated patient-derived ECM from three donors with different causal mutations. We found that the structure, content, and mechanics of the ECM depended not only on whether a patient has vEDS, but also on the specifics of the driving mutations. These results demonstrate a connection between the biomechanical response of ECM and *COL3A1* mutation, in addition to differences in protein composition other than collagen III, including glycosaminoglycans. The resulting biochemical and biomechanical characterization of patient cell-derived ECM (CDM) enables further mechanistic studies and potential therapeutic interventions to better understand and treat vEDS based on the specific causal genetic mutations. Further, the development of methods to rigorously characterize ECM from donor fibroblasts is likely to have broad applicability in diseases in which ECM composition and mechanics play a role in progression.

## 2. Materials and Methods

### 2.1 Cell culture

Human dermal fibroblasts were obtained from either the Coriell Institute of Medical Research biobank or ATCC. Information regarding age, sex, mutation, and source information can be found in Table S1. Fibroblasts were cultured at 37°C, 5% CO_2_ in DMEM (Gibco 11885-844) + 10% vol/vol FBS (Avantar 97068-085) and used between passage 5 and 25. Human aortic endothelial cells (HAECs) were cultured at 37°C, 5% CO_2_ in EGM-2 Bullet Kit (Lonza CC-3162) medium and used between passages 4-8.

### 2.2 Fibroblast characterization

For characterization, fibroblasts were plated at 1,750 cells/cm^2^ on 35 mm glass-bottom dishes (Mattek P35GC-1.0-14-C) coated with 0.2% (m/v, in PBS) gelatin (Sigma G1393). After 24 hours, fibroblasts were fixed with 4% (w/v, in PBS) paraformaldehyde (Electron Microscopy Sciences 15710), permeabilized with 0.1% (w/v, in DI water) Triton X-100 (Sigma 1003339792) for 20 minutes at RT and stained with AlexaFluor-488 conjugated phalloidin (Invitrogen A12379, diluted 1:200) and DAPI (Fisher 62248, diluted 1:1000). Fibroblasts were then imaged on an Olympus IX83 Inverted Microscope with a 10X/NA 0.4 objective. For quantification, images were run through a custom pipeline in CellProfiler to assess cell shape characteristics of area, perimeter, eccentricity, and form factor [28].

### 2.3 Gene and protein expression

Fibroblasts were plated at a density of 26,000 cells/cm^2^ and allowed to recover from passaging for 24 hours before extracting RNA using TRIZOL (Invitrogen). RNA was recovered through precipitation with chloroform and isopropanol. RNA concentration was measured using a SpectraMax QuickDrop (Molecular Devices), prior to conversion to cDNA using the Superscript IV VILO with ezDNase kit (ThermoFisher #11766050), and a MaxyGene II thermal cycler (Corning). Real-time PCR was performed in 5 μL reactions using the PowerUp SYBR Green Mastermix (ThermoFisher #A25742) and a QuantStudioI 6 Flex System (ThermoFisher). Specifics regarding cycle parameters and primers can be found in Supplementary Methods. For protein expression levels, fibroblasts were plated at 26,000 cells/cm^2^ in 6-well plates. Media was changed 24 hours after plating and either included 50 µg/mL ascorbic acid (+AA, Sigma A5960) or did not include ascorbic acid (-AA). Cell lysate was collected 24 hours after media change by lysing cells with RIPA Buffer (ThermoFisher, 89901) and HALT protease and phosphatase inhibitor cocktail (ThermoFisher, 1861280) for 5 minutes in the dish. Then, the dishes were scraped, and the lysate was transferred to a microcentrifuge tube. Samples were incubated for another 5 minutes and then sonicated 3 times at an amplitude of 30% in 10 second cycles using a Branson 150 Sonifier. Protein concentration in total cell lysate was calculated using the BCA Gold Assay (ThermoFisher, A53226). After protein concentration was determined, samples were mixed with Bold Sample Reducing Agent (Invitrogen, B0004) and NuPAGE LDS Sample Buffer (Invitrogen, NP0007), then were boiled for 5 minutes in a MaxyGene II thermal cycler (Corning). Samples were stored at -80 °C prior to gel electrophoresis. Details regarding Western Blot procedures and antibodies are presented in the Supplementary Methods.

### 2.4 ECM production

ECM production was monitored using confocal reflectance microscopy on an Olympus FV3000 laser scanning confocal microscope with a 488 nm laser and 60x/NA 1.4 objective. Images were taken at 1-µm z-step, and the thickness of each sample was determined by extracting the z-axis intensity profiles in FIJI and then using a custom MATLAB script to determine the difference between the most intense z-plane and the first z-plane below the cut off value corresponding to that above the surface (determined to be 500 AU).

### 2.5 Traction force microscopy

Traction force microscopy was carried out according to previous protocols [29-31]. Briefly, 8 kPa polyacrylamide (PAA) gels were prepared on glass bottom dishes (Mattek P35GC-1.0-14-C) using a coverslip to localize 0.1 µm (580/605) FluoSphere carboxylated fluorescent beads (Invitrogen F8801) to the gel surface. After curing, the glass coverslip used to localize the beads was removed, and a 0.2% (m/v, in PBS) gelatin (Sigma G1393) coating was conjugated to the polyacrylamide gels using *N-*ethyl-*N’*-(3-(dimethylamino)propyl)carbodiimide/*N*-hydroxysuccinimide (EDC/NHS) chemistry. Briefly, PAA gels were incubated for 1 hour at room temperature with a solution of 137 mM NaCl and 5% (w/v, in DI water) glycerol and then incubated with an EDC/NHS buffer comprised of DI water, 10X EDC (19 mg/mL in DI water), 10X NHS (29 mg/mL in DI water), and 2X conjugation buffer of 0.2 M 2-ethanesulfonic acid (MES) and 10% (w/v, in DI water) glycerol at pH 4.5 for 30 minutes at room temperature. Subsequently, PAA gels were incubated with 0.2% (m/v, in PBS) gelatin for 1 hour at 37 °C and stored in PBS at 4 °C prior to use. Fibroblasts were then plated onto the PAA gels at a density of 1,593 cells/cm^2^, and after 24 hours, traction force microscopy was carried out using a Nikon Eclipse inverted Widefield microscope with an Oko labs live-cell environmental chamber using a 20X/NA 0.75 objective with a 1.5X tube lens. Images of fibroblasts were obtained every five minutes for 30 minutes before 100 µL of a 5% w/w solution of SDS (sodium lauryl sulfate, Sigma L3771) was added to remove fibroblasts. Imaging was continued for 30 minutes to collect a reference state image for traction force analysis. Details on image analysis can be found in the Supplementary Methods.

### 2.6 Cell sheet contraction

Cell sheets were generated by plating fibroblasts on gelatin-coated 48-well plates at a density of 26,000 cells/cm^2^. Media was supplemented with 50 µg/mL ascorbic acid (Sigma A5960) and changed daily for 30 days. For quantification, cell sheets were imaged daily starting on day 3 with an Olympus IX83 inverted widefield microscope using phase contrast with a 4X/NA 0.16 objective (**Fig. S1**). Images were stitched using the Stitching plug-in on FIJI [32], then the outline of the cell sheet was drawn using the polygon tool and the enclosed area was recorded. The size of the well was recorded using the ellipse tool, and the cell sheet area was normalized to the well area for each image. To visualize cell distribution, cells were stained with CellMask Deep Red (Invitrogen, C10046) on day 2 and after full contraction. After staining, cell sheets were imaged with an Olympus IX83 inverted widefield microscope and for visualization images were stitched using the Stitching plug-in on FIJI [32].

### 2.7 Collagen gel contraction assay

To house collagen hydrogels, 4 mm diameter PDMS wells were passively adhered to #1 9 mm x 9 mm glass coverslips (Electron Microscopy Sciences #72190-09). Wells were passivated overnight using vapor deposition of Trichloro(1H,1H,2H,2H-perfluorooctyl)silane (Sigma 448931) and sterilized with 70% (v/v) ethanol in DI-H_2_O followed by UV sterilization for 15 min before placing into a 24-well plate for cell culture. Collagen type I derived from rat tail (Corning 354326) was buffered with 10x DMEM and 10x reconstitution buffer (0.3 M NaHCO_3_ and 0.4 M HEPES in DI-H_2_O), titrated to a pH of 8.0 with NaOH, and brought to a final concentration of 2 mg/mL. Fibroblasts were suspended at a concentration of 40,000 cells/mL of collagen precursor solution, and the solution was added to each PDMS well and allowed to polymerize for 30 minutes at 37°C. Media was then added to hydrate the hydrogels, and each hydrogel was imaged daily on an Olympus IX83 inverted microscope at 4x magnification. Images were stitched using the Stitching plug-in on FIJI [32] and the enclosed area was recorded using the polygon tool. The area was normalized to the day 1 measurement for each sample.

### 2.8 Cell derived matrix generation and biochemical characterization

Cell-derived matrix (CDM) was generated with slight modifications to a previously used protocol [33]. Briefly, Fibroblasts were plated at 26,000 cells/cm^2^ on dishes (35 mm Mattek dishes or well plates) coated with 0.2%(m/v) gelatin. Fibroblasts were treated with ascorbic acid (50 µg/mL, Sigma A5960) added to the growth medium for 6 days. After 7 days, samples were decellularized using an extraction buffer of 50 mM NH_4_OH and 0.5% Triton X-100 in PBS to remove fibroblasts, followed by a DNAse I (10 µg/mL, Sigma D4527) treatment to remove residual DNA. Decellularization was confirmed by staining ECM with AlexaFluor-488 conjugated phalloidin (Invitrogen A12379) and DAPI (Fisher 62248). After decellularization, CDM was stored in PBS + 1x Pen/Strep (Gibco 15140-122) + 0.25 µg/mL Amphotericin B (Sigma A2942) until analysis and used within 4 weeks from decellularization. Decellularized CDM was digested prior to proteomic characterization (Supplemental Methods).

### 2.9 Scanning Electron Microscopy (SEM) imaging and quantification

CDM was generated on 9 × 9 mm coverslips in 6-well plates as described above and was serially dehydrated from 30% (v/v, in water) ethanol to 100% (v/v, in water) ethanol over 24 hours, increasing ethanol concentration every hour, and once reaching 100% incubating overnight. Samples were then dried with critical point drying using a Tousimis 931.GL critical point drier. Dried samples were then transferred to sample holders and sputter coated with 8 nm of Au-Pd. SEM imaging was then carried out on a Hitachi S-4700 scanning electron microscope. Five images were taken at random locations per sample at 25,000X magnification. For quantification, SEM images were segmented using the machine learning based Weka segmentation plugin [34] for FIJI [35], which uses pixel-wise binary classification. The classifiers were trained by manually tracing along fibril or the background and classifying the traced lines as class 1 (indicating fibril) or class 2 (indicating background) until the trained model effectively identified fibrils in the segmented images. Segmented images were input into DiameterJ [36] to quantify fibril size and pore area.

### 2.10 Collagen and GAG content

CDM was generated in 12-well plates and stained with Picrosirius Red (Sirius Red/Direct Red 80, Sigma 36554) for 1 h at RT on an orbital rocker. Stained samples were then washed for 30 min at RT with 0.01 N Hydrochloric acid and imaged using an Olympus BX60F5 microscope with a 10X/NA 0.30 objective with an Olympus DP73 color camera. After imaging, the stain was released to determine relative collagen concentration [37] by incubating with 0.1 N NaOH for 30 minutes on an orbital rocker at RT, collecting supernatant from each well, and measuring absorbance at 550 nm in a plate reader against a 0.1 N NaOH blank. Absorbance values from the blank were then subtracted from the Picrosirius Red stain absorbance values. GAG content was quantified using a Glycosaminoglycans Assay Kit (Chondrex, inc. #6022) following manufacture specifications. Briefly, CDM was generated in 12-well plates and solubilized with 125 µg/mL papain by scraping the CDM into a tube and incubating with papain for 3 h at 65 °C with occasional mixing. After solubilization, tubes were incubated at 90 °C for 10 min to inactivate the papain and centrifuged at 10,000 RPM for 10 minutes at RT and supernatant was collected. 50 µL of the solubilized sample was added to the well of the kit-provided clear-bottom well plate along with 50 µL of 1,9 Dimethylmethlyene Blue (DMB) dye (or PBS as a negative control) was added to each well to stain sample. Absorbance values at 525 nm were measured within 5 min of adding DMB using a plate reader. For quantification, PBS control sample absorbances were subtracted from stained sample absorbances. Then these absorbance values were normalized to the absorbance of the first sample loaded on the plate on which it was run to determine relative differences in GAG concentration between samples.

### 2.11 Mechanical analysis

CDM was generated in 35 mm glass-bottom dishes as described above, and mechanical measurements were taken using a Piuma nanoindentor (Optics11) equipped with a 0.030 N/m cantilever with a 9 µm diameter spherical tip. Samples were indented at a loading rate of 300 nm/s at 25 different points on each sample in indentation mode (**Fig. S2**). Data were exported and fit to a modified Hertzian contact model to account for the thin sample (Supplemental Methods) using a series of custom MATLAB scripts. For stress relaxation studies, CDM was indented at a loading rate of 3000 nm/s to an indentation depth of 300 nm, which was maintained for five seconds before unloading (**Fig. S2**). Stress relaxation curves were collected for 25 different points on each sample, and data were exported and analyzed using a series of custom MATLAB scripts (Supplemental Methods).

### 2.12 Human arterial endothelial cell (HAEC) Migration Analysis

CDM was generated in 35 mm glass-bottom dishes as described above, and HAECs (Lonza) were plated onto CDM at a density of 12,500 cells/cm^2^. HAECs were allowed to recover from passaging for 24 h after plating and were subsequently incubated with AlexaFluor-647 anti-human VE-Cadherin antibody (1:100, BD Biosciences, #561567). After 15 minutes incubation, HAECs were imaged live for a period of 7 h taking an image every 35 s using an Olympus FV3000 laser scanning confocal microscope with a 640 nm laser and 30x U Plan S-Apo N 1.05 NA silicone oil immersion objective. Images were analyzed using the Cellpose TrackMate plug-in in FIJI [38] to extract metrics related to the migration of the HAECs including median migration speed, mean straight line speed (net distance/total track time), and persistence (net distance/total distance traveled).

### 2.13 Statistics

Young’s moduli data were statistically assessed by first applying a log transform and then using a linear mixed model or robust linear mixed model in R [39] with p < 0.05 indicating significance. For the robust linear mixed model, the degrees of freedom were assumed to be the same as the traditional linear mixed model for determining p-values of statistical comparisons. All other statistical analyses were run with Prism 9.0 with p < 0.05 indicating significance. Unpaired student’s t-test, one-way ANOVA, or two-way ANOVA were run where appropriate as indicated in figure captions with a Tukey post-hoc test between all conditions.

## 3. Results

### 3.1 vEDS patient-derived fibroblasts

Dermal fibroblasts from healthy and vEDS patient donors were obtained from the NIGMS Biobank through the Coriell Institute and used throughout this study. Three donors from patients diagnosed with vEDS were acquired, including one patient with an in-frame base pair change (*COL3A1*^+/IVS40-1G>A^), one patient with a frameshift mutation (*COL3A1*^+/766delA^), and one patient with a glycine substitution variant (*COL3A1*^+/G939D^). The molecular characterization was performed before cell line submission to Coriell or by clinical summary/case study as indicated in the Coriell Institute National Institute of General Medical Sciences (NIGMS) biobank (**Table S1**). Commercially available healthy neonatal fibroblasts were used as a comparison for healthy expression of *COL3A1*. vEDS patient-derived fibroblasts showed differential magnitude of *COL3A1* transcript expression dependent on donor genotype (**Fig. 1a**). Fibroblasts harboring *COL3A1*^+/IVS40-1G>A^ and *COL3A1*^+/766delA^ mutations expressed fewer *COL3A1* transcripts, as opposed to fibroblasts harboring *COL3A1*^+/G939D^ mutations, which demonstrated increased levels of *COL3A1* compared to healthy neonatal fibroblasts (**Fig. 1a**). Interestingly, fibroblasts from vEDS patients demonstrated decreased expression of COL3A1 protein across all donor genotypes, a finding that persisted when treated with ascorbic acid to promote collagen deposition (**Fig. 1b-c**). We further investigated variation in cell shape and cytoskeletal morphology in vEDS patients and healthy fibroblasts. We plated patient-derived and healthy fibroblasts at a low density and observed that vEDS patient-derived fibroblasts were significantly larger in area than healthy fibroblasts and displayed variable shape, as measured by cell eccentricity, with cells expressing the glycine substitution mutation characterized by significantly lower eccentricity (**Fig. 1d-e, Fig. S3**).

**Fig. 1.**
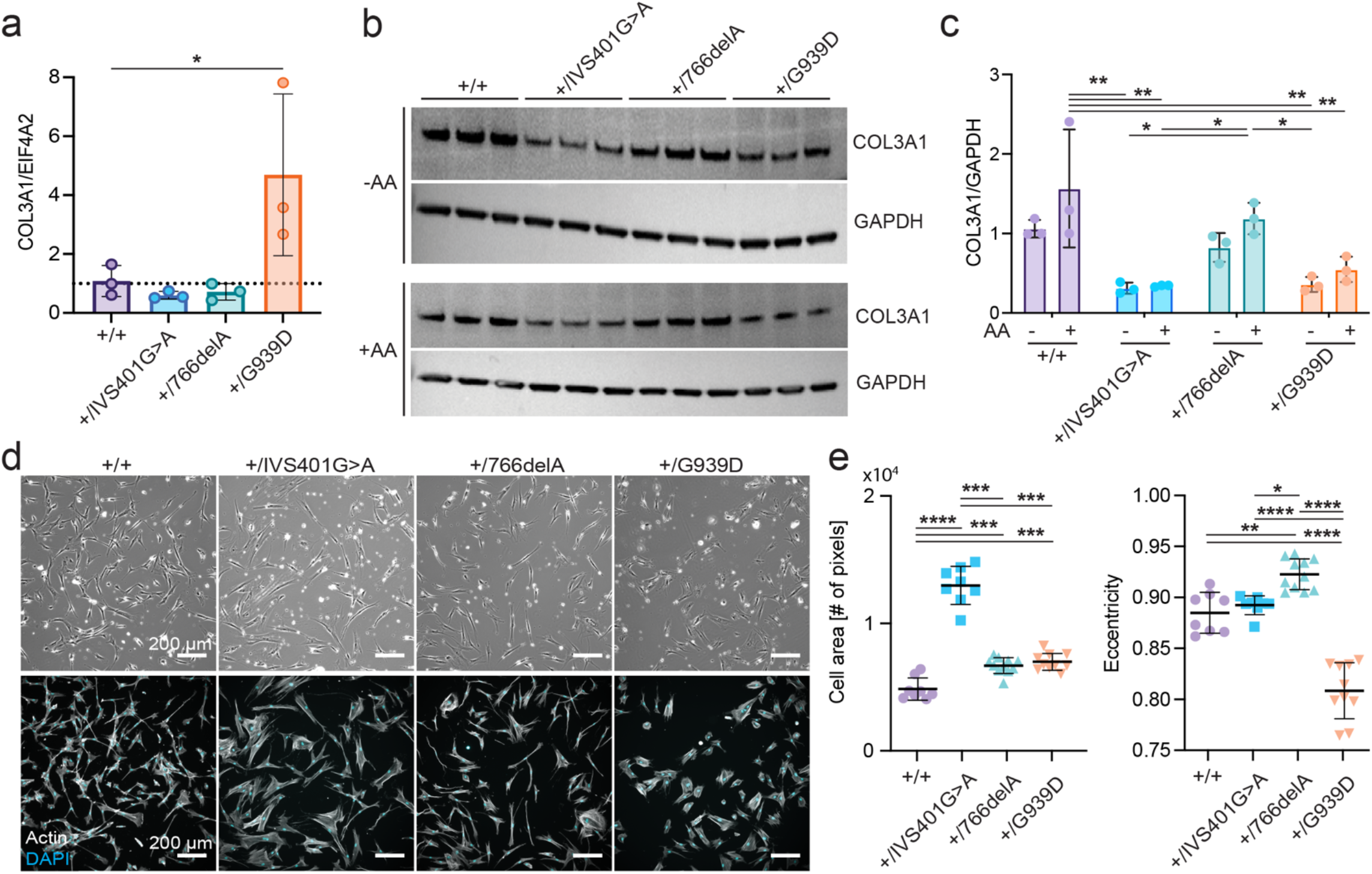
COL3A1 mutant fibroblasts demonstrate differential COL3A1 expression and cell shape. **a** Relative gene expression of *COL3A1* measured by qRT-PCR in cultured fibroblasts (*n* = 3 independent lysates, statistics shown are a one-way ANOVA with a Tukey post-hoc test). **b** Expression of COL3A1 protein as measured by western blot from fibroblasts cultured with and without ascorbic acid (AA) treatment. Each lane represents an independent lysate. **c** Quantification of western blot band intensity (*n* = 3 independent lysates, statistics shown are a two-way ANOVA with a Tukey post-hoc test). **d** Fluorescence micrographs of fibroblasts stained with DAPI (nucleus) and phalloidin (F-actin). **e** Area and eccentricity of cells, quantified from phalloidin-stained fibroblasts. Each data point is the average area or eccentricity of all cells in an image from an independently prepared sample (*n* > 8 independent samples, statistics shown are a one-way ANOVA with a Tukey post-hoc test). All plots, mean ± standard deviation, *P<0.05, **P<0.01, ***P<0.001, ****P<0.0001.

### 3.2 ECM synthesis

To determine whether patient genotype impacts ECM assembly and deposition, we supplemented the culture media with ascorbic acid to stimulate ECM synthesis [33, 40]. We used confocal reflectance microscopy to measure matrix structure and deposition non-destructively over time by illuminating samples with 488 nm light, which has been shown to be capable of resolving fibers with diameters greater than 250 nm [41-43] (**Fig. 2a-b, Fig. S4a**). Interestingly, we found that the rate of ECM deposition was a function of patient genotype, and fibroblasts exhibiting *COL3A1*^+/IVS40-1G>A^ and *COL3A1*^+/766delA^ mutations deposited significantly thinner layers of ECM than fibroblasts harboring *COL3A1*^+/G939D^ mutations, which deposited layers of similar thickness to the neonatal healthy control (**Fig. 2b**).

**Fig. 2.**
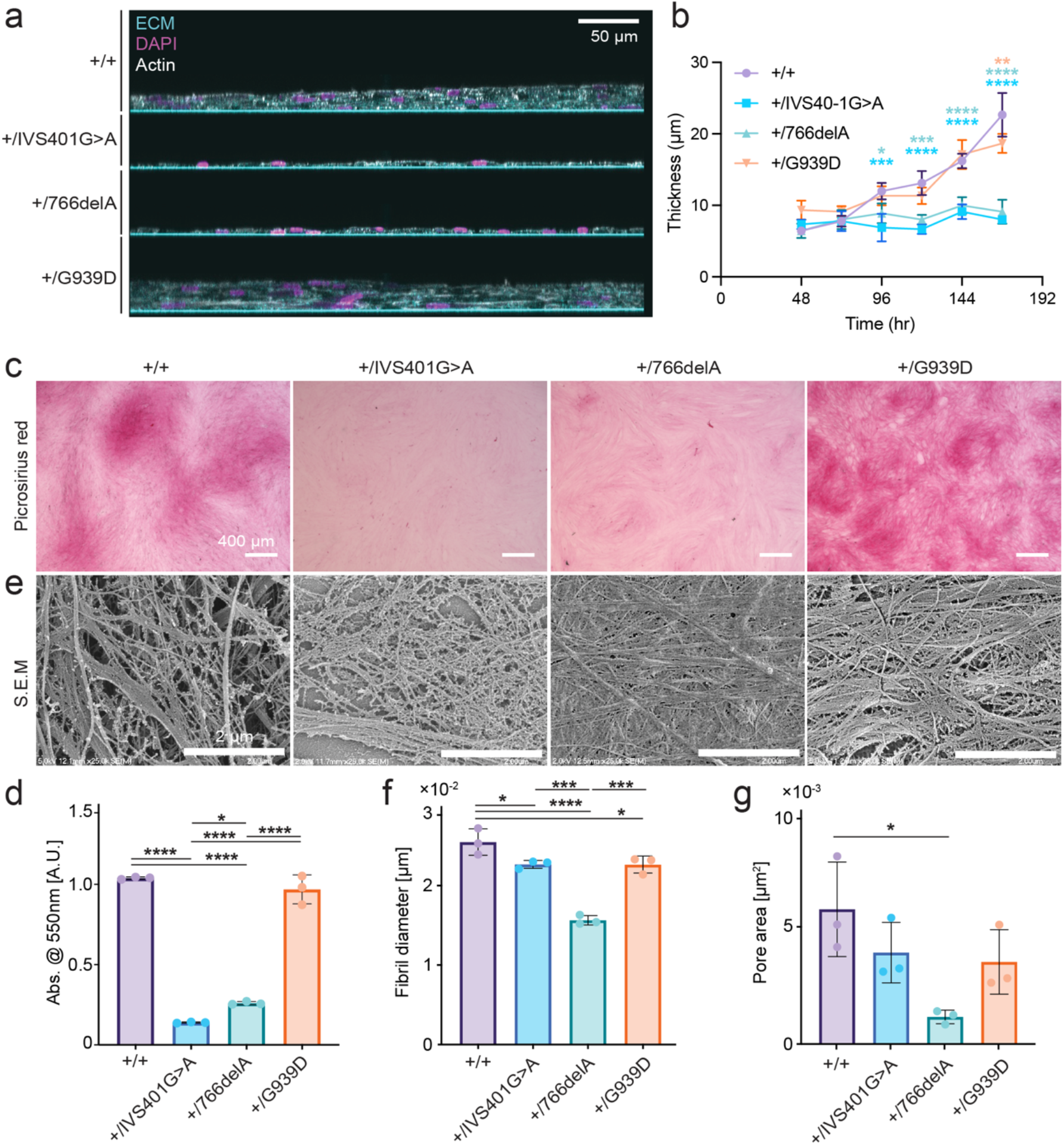
ECM assembly is altered in vEDS fibroblasts. **a** Composite x-z projection micrographs of fibroblasts and deposited ECM stained with phalloidin (F-actin) and DAPI (nucleus). ECM was visualized with confocal reflectance microscopy at 488 nm (scale bar = 50 µm). **b** Thickness of deposited ECM as measured daily by confocal reflectance microscopy at 488 nm (line indicates mean and corridor indicates standard deviation, as measured from *n* = 3). Statistics shown are indicative of the statistical significance between +/+ and +/IVS40-1G>A (blue), +/+ and +/766delA (green) or +/+ and +/G939D (orange). The statistical test performed was a two-way ANOVA with a Tukey post-hoc test (*P<0.05, **P<0.01, ***P<0.001, ****P<0.0001) **c** Collagen content was assessed with picrosirius red staining, and **d** collagen content was quantified by stripping bound picrosirius red stain and measuring absorbance at 550 nm. **e** ECM fibril structure was assessed with scanning electron microscopy (S.E.M.). **f-g** Fibril diameter and inter-fibril pore area was measured from S.E.M. micrographs (n = 3 images). Statistics shown are a one-way ANOVA with a Tukey post-hoc test (*P<0.05, **P<0.01, ***P<0.001, ****P<0.0001), all plots are mean ± standard deviation, and micrographs are representative of images taken from at least 3 independent samples.

To further probe the structure and content of ECM deposited by donor cells, we removed cellular content from the deposited ECM and verified decellularization by staining for filamentous actin and DNA after treatment with extraction buffer (**Fig. S4b-c**).We then assessed collagen content of the resulting cell-derived matrix (CDM) using picrosirius red (PSR) staining and scanning electron microscopy (S.E.M.). Micrographs of PSR staining and quantitative measurement of PSR bound to patient CDM demonstrated significantly lower collagen content in CDM derived from fibroblasts harboring *COL3A1*^*+/IVS40-1G>A*^ and *COL3A1*^*+/766delA*^ mutations compared to CDM from fibroblasts harboring *COL3A1*^*+/G939D*^ mutations and the neonatal healthy fibroblasts (**Fig. 2c-d**). The structure of the CDM at the fibril level also varied with patient genotype (**Fig. 2e**), with the average fibril diameter measuring significantly smaller in CDM from all vEDS donors compared to healthy CDM (**Fig. 2f**) and pore area trending smaller in patient CDM, though only statistically significant for CDM from the *COL3A1*^*+/766delA*^ donor.

### 3.3 ECM Proteomic Analysis

While the *COL3A1* mutations expressed by all donors are associated with increased incidence of aneurysm formation and rupture, patients harboring the glycine substitution mutation (*COL3A1*^*+/G939D*^) are at a significantly higher risk for aneurysm and present clinically with more severe disease characteristics [44, 45]. To study the impact of specific causal mutation on the biochemical composition of deposited ECM, we conducted unbiased proteomic characterization of constituent proteins in CDM derived from each donor and the healthy fibroblast control. We solubilized samples using a combination of elastase and trypsin and used label-free LC-MS quantification methods to determine peptide sequences present in each sample. We then determined the protein expression profiles for which proteins were significantly up-/down-regulated between each patient donor CDM and healthy CDM (**Fig 3, S5**).

**Fig. 3.**
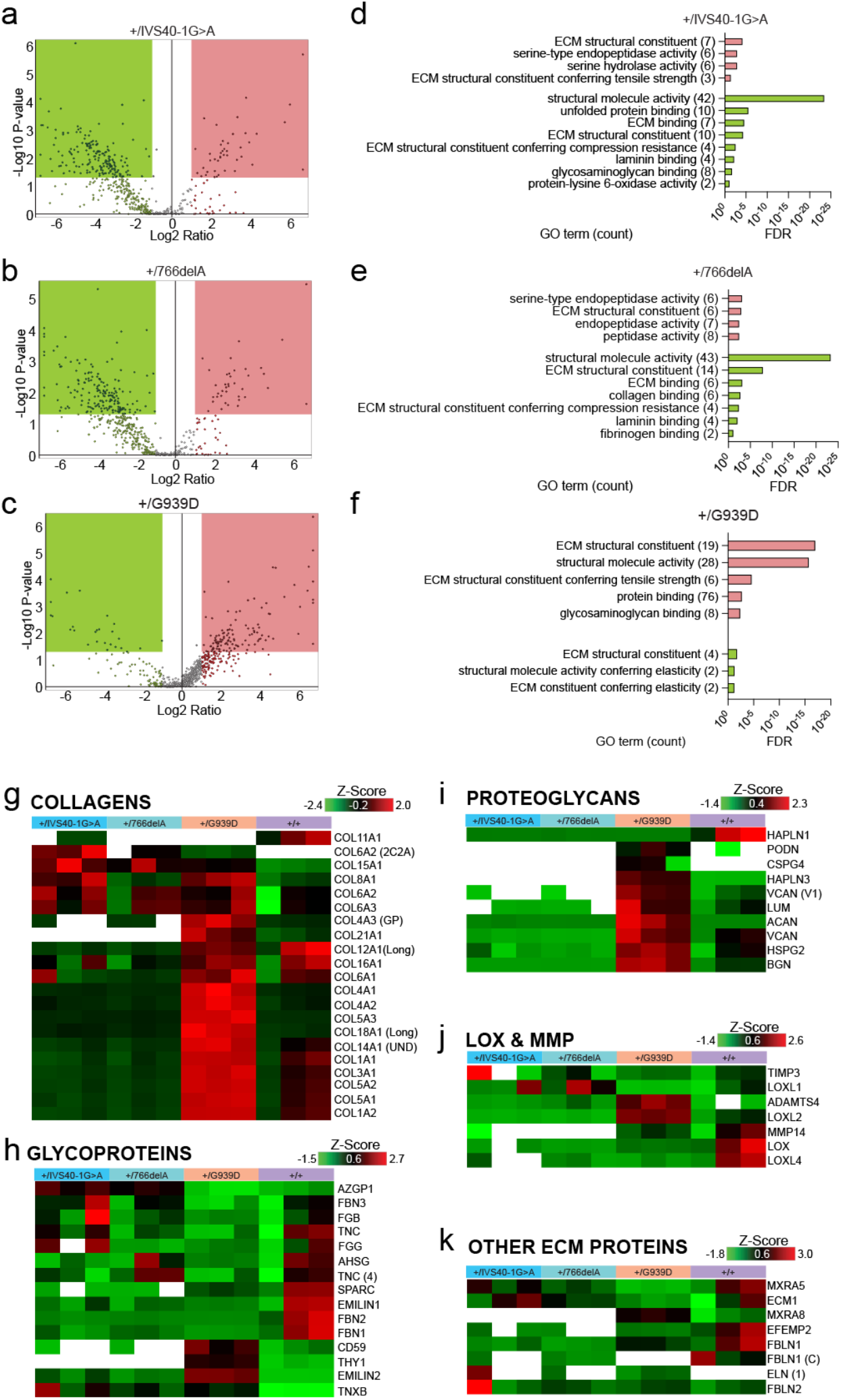
ECM proteomic characterization reveals differences in ECM composition. **a-c** Volcano plot visualizing upregulated and down regulated proteins for +/mutant CDM each compared to +/+ CDM. To determine significant up- (red box) and down-regulation (green box), cut off values of X = log_2_(2) and Y = - log_10_(0.05) were used. **d-f** Sampled results of GO analysis for molecular function of significantly up- or downregulated proteins for +/mutant CDM each compared to +/+ CDM, presented as false discovery rate (FDR). Down regulated GO terms are shown in green (bottom) and upregulated GO terms are shown in red (top), and the number of proteins associated with each GO term is shown in parentheses. **g-k** Heat maps of proteomic characterization of collagens, glycoproteins, proteoglycans, Lysyl-oxidases, MMPs, and other associated ECM proteins. These maps show normalized abundance values for each replicate (n = 3 replicate per genotype) scaled to a z-score before hierarchical clustering using the Euclidian distance function and the complete linkage method. White spaces indicate that the protein was not found in that sample.

We observed a divergence in protein content depending on genotype when we looked at significant up- and downregulation (**Fig 3a-c**). The CDM generated from fibroblasts with the base pair substitution and frameshift (*COL3A1*^*+/IVS40-1G>A*^ and *COL3A1*^*+/766delA*^, respectively) had a larger number of significantly downregulated proteins (176 and 181 proteins) compared to healthy (**Fig. 3a**,**b**), whereas CDM generated from fibroblasts with the glycine substitution mutation (*COL3A1*^*+/G939D*^) had a larger number of significantly upregulated proteins (120 proteins) (**Fig 3c**). To determine the function of these differentially expressed proteins, we performed gene ontology (GO) term enrichment analysis for molecular function (**Fig. 3d-f**). Interestingly, we found that in all patient donor CDM, the GO terms with the highest count for differential protein expression related to ECM structure, structural integrity, and mechanics (**Fig. 3d-f**).

We then sought to identify the individual proteins that constitute the differential GO term enrichment by investigating abundance levels of individual ECM-related proteins. The abundance of collagen (**Fig 3g**) was consistent with the results of PSR staining (**Fig 2c**,**d**), and the decreased (*COL3A1*^*+/IVS40-1G>A*^ and *COL3A1*^*+/766delA*^) or increased (*COL3A1*^*+/G939D*^) abundance of COL3A1 as compared to healthy (**Fig 3g**) was consistent with the gene expression and protein levels in cultured fibroblasts determined by PCR and Western blot, respectively (**Fig 1a**,**b**). Interestingly, COL6A2, COL6A3, and COL15A1 were upregulated in all *COL3A1*^*+/mutant*^ CDM compared to healthy control (**Fig 3g**). A subset of glycoproteins, including SPARC, EMILIN1, FBN2, and FBN1, were present at lower abundance in all *COL3A1*^*+/mutant*^ CDM compared to healthy control (**Fig 3h**), while TNXB and EMILIN2 were present in higher abundance in all mutant CDM compared to healthy control (**Fig 3h**). In contrast, proteoglycans were more distinctly correlated with genotype. HAPLN1 was the only protein with consistent decrease in abundance among all *COL3A1*^*+/mutant*^ CDM as compared to the healthy control (**Fig 3i**). All other proteoglycans, including ACAN, BCAN, LUM, and VCAN showed increased abundance in *COL3A1*^*+/G939D*^ CDM compared to healthy control and compared to other COL3A1^+/mutant^ CDM (**Fig 3i**). In addition to proteoglycans, matrix-modulating proteins MMP14, LOX, and LOXL4 were found in higher abundance in all *COL3A1*^*+/mutant*^ CDM compared to healthy control (**Fig 3j**), while ADAMTS4 and LOXL2 were increased only in COL3A1^+/G939D^ CDM (**Fig 3j**). While not changed in *COL3A1*^*+/G939D*^ CDM, TIMP3 and LOXL1 were found in increased abundance in *COL3A1*^*+/IVS40-1G>A*^ and *COL3A1*^*+/766delA*^ CDM (**Fig 3j**). Of the other examined ECM proteins, FBLN1 and EFEMP2 were decreased in all *COL3A1*^*+/mutant*^ CDM compared to healthy control (**Fig 3k**), while MXRA5 and ECM1 were decreased in only *COL3A1*^*+/G939D*^ CDM compared to healthy CDM (**Fig 3k**). Interestingly, isoform 1 of elastin was only found in *COL3A1*^*+/mutant*^ CDM (**Fig 3k**). Collectively, these results demonstrate complex changes to the matrix milieu in vEDS patients and suggest that the biochemical mechanisms leading to increased aneurysm formation could vary depending on patient genotype.

### 3.4 Tissue contraction

To determine the functional consequences of the differences in biochemical composition and structure of CDM from healthy and vEDS patient donors, we plated fibroblasts at confluence in wells of a 48-well plate and treated with ascorbic acid for 30 days. We imaged the cell-laden ECM each day (**Fig 4a, Fig. S1a**) and over time, cells deposited and eventually contracted ECM (**Fig 4a, Fig. S1**). The time to contraction varied significantly depending on the fibroblast genotype (**Fig. 4b**) and inversely correlated with collagen content, with *COL3A1*^*+/G939D*^ and healthy fibroblast-ECM constructs contracting within 15 days, and *COL3A1*^*+/IVS40-1G>A*^ and *COL3A1*^*+/766delA*^ fibroblast-ECM constructs contracting around 30 days (**Fig. 4b**).

**Fig. 4.**
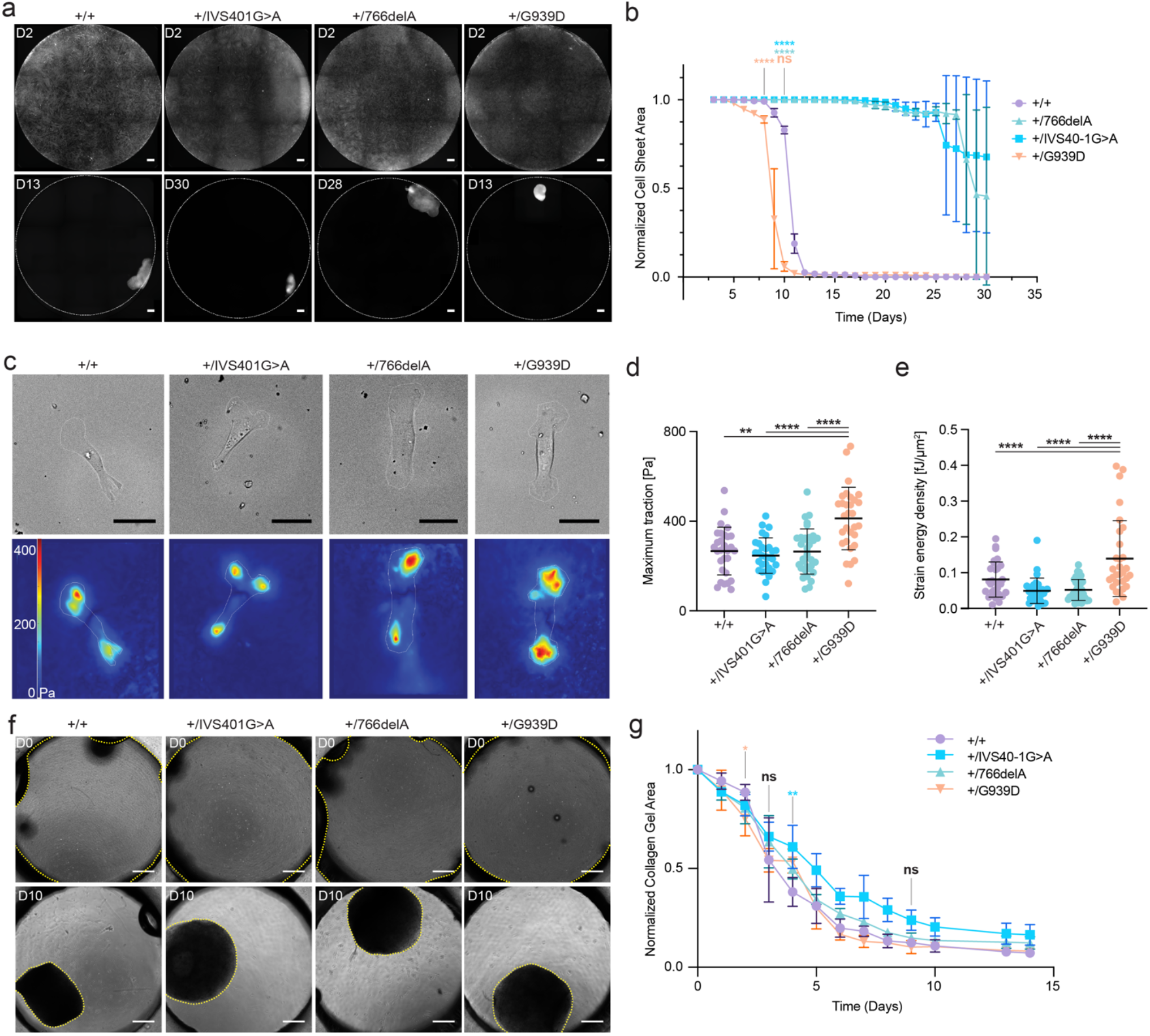
Cell-generated ECM contraction depends on genotype. **a** Cell sheet contraction visualized with fluorescent cytoplasmic stain on day 2 after plating (top) and on the day of contraction (bottom, scale bar = 500 µm). **b** Cell sheet area normalized to the area of the well as a function of time. **c** Brightfield images of cells (top) and traction maps (bottom) as measured by TFM of COL3A1^+/+^ and COL3A1^+/mutant^ fibroblasts on gelatin-coated 8 kPa PAA gels (scale bar = 50 µm). **d** Maximum traction stress and **e** strain energy density calculated from traction maps. Statistics shown from one-way ANOVA with a Tukey post-hoc test. **f** COL3A1^+/+^ and COL3A1^+/mutant^ fibroblasts embedded within 2 mg/mL collagen I hydrogels after seeding (top) and on day 10 (bottom, yellow dotted line indicates perimeter of cell-laden hydrogel, scale bar = 500 µm). **g** Cell-laden collagen hydrogel area normalized to the area on day 0. For all subpanels, n ≥3 for each condition, plots are mean +/- standard deviation. *P<0.05, **P<0.01, ***P<0.001, ****P<0.0001 as determined by two-way ANOVA with a Tukey post-hoc test, and comparisons between genotypes are indicated by color, with blue indicating comparison between +/+ and +/IVS40-1G>A, green indicating +/+ vs. +/766delA, orange indicating +/+ vs. +/G939D. Black indicates that a statistically significant difference when compared to all other conditions).

To determine whether the observed differences in ECM contraction were due to differences in fibroblast contractility, we performed traction force microscopy (TFM) [29-31] on isolated cells cultured on 8 kPa polyacrylamide hydrogels functionalized with gelatin (**Fig. 4c**). We found that *COL3A1*^*+/G939D*^ fibroblasts exerted larger maximum traction magnitudes (**Fig. 4d**) and had a larger strain energy density (**Fig. 4e**) compared to other cell types, which were similar in magnitude to the healthy control cells. To further explore whether there were differences in fibroblast contractility when cells were embedded in a standardized 3D ECM, we suspended fibroblasts in reconstituted 2 mg/mL type I collagen hydrogels (**Fig. 4f**). We measured the area of cell-laden hydrogels over 14 days and unlike the cell-sheet contraction (**Fig. 4b**), the rate of collagen hydrogel contraction did not depend on fibroblast genotype (**Fig. 4g**). Collectively, these results suggest that the observed differences in the contraction rate of cell-laden, cell-generated CDM derived from patient cells are not due to differences in baseline contractility of the fibroblasts themselves.

### 3.5 ECM elastic mechanical properties

We hypothesized that differences in mechanical properties of cell-generated ECM were responsible for the observed differences in rates of contraction of cell sheets as a function of donor genotype (**Fig. 4a-b**). To test this hypothesis, we synthesized cell-generated ECM as above, and prior to ECM contraction, we removed cells to generate acellular patient donor CDM (**Fig. S4**). We then characterized the mechanical properties of the CDM using nanoindentation. To determine elastic mechanical properties, we indented each sample with a 0.030 N/m stiffness probe with a 9 µm diameter spherical tip to an indentation depth of 300 nm at a loading rate of 300 nm/s. Each sample was indented at 25 different points in a 5×5 grid with 100 µm linear spacing between each indentation point (**Fig. S2**). We analyzed indentation vs. load curves (**Fig. 5a**,**b**, **S7a**) using a modified Hertzian-contact fit model that accounts for the finite thickness of the CDM samples [46].

**Fig. 5.**
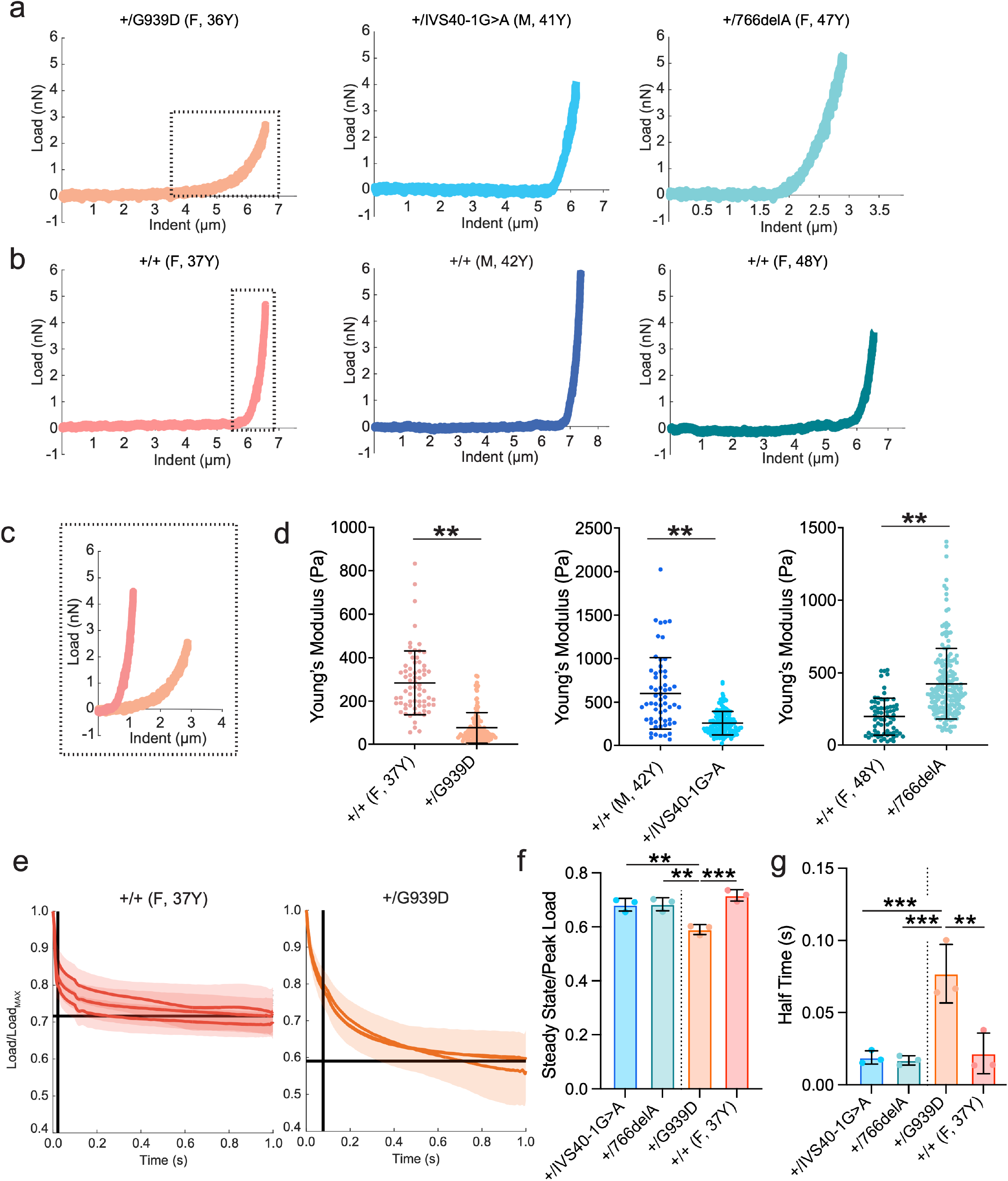
CDM elastic and time-dependent mechanical properties are a function of *COL3A1* mutations. **a-b** Representative load v. indentation curves for +/mutant CDM and age and sex-matched healthy control CDM. **c** Overlay of region indicated by dotted black lines for +/G939D vs. age/sex matched +/+ CDM. **d** Young’s modulus of CDM generated from the +/G939D, +/IVS40-1G>A, and +/766delA donors compared to CDM from their age and sex-matched healthy control donor (statistical significance determined by a linear mixed model). **e** Stress relaxation (load vs. time) curves for CDM generated from the +/G939D donor and an age/sex-matched donor (+/+ F, 37Y). Corridor shows standard deviation from mean (solid colored line) for each sample. Black horizontal lines indicate the average steady-state/peak stress for all samples. Black vertical lines indicate the average half time for all samples. **f** Steady state load normalized by the steady state load measured 1s after indentation, and **g** half-time of relaxation. (*p<0.05, **p<0.01, ***p<0.001, ****p<0.0001 for all panels as determined by one-way ANOVA with Tukey post-hoc test unless otherwise indicated. For all data, n ≥ 3 biological samples, data points represent individual Young’s modulus value, and data are presented as mean +/- standard deviation).

ECM mechanical properties have been shown to change with age [47], and the vEDS donor fibroblasts were derived from patients with various age and sex demographics (**Table S1**). To account for difference in donor age, we acquired fibroblasts from three healthy donors with similar age/sex to each patient donor in addition to our initial healthy fibroblast line (**Table S1**). These fibroblasts expressed similar levels of *COL3A1* (**Fig. S6a**) and generated ECM at a slightly reduced rate compared to the neonatal healthy control (**Fig. S6b**). Using a robust linear mixed statistical model (**Table S5**), we found that the Young’s modulus varied significantly with age (**Fig. S7c**) and sex (**Fig. S7d**). Interestingly, *COL3A1*^*+/G939D*^ CDM was significantly more compliant than the age and sex matched healthy control (**Fig. 5d**). Notably, the magnitude of Young’s modulus correlated with the rate of cell sheet contraction (**Fig. 5c, Fig. 4b**), further suggesting that changes in mechanical properties of the ECM drive the observed differences in rates of contraction.

### 3.6 ECM time-dependent mechanical properties

Given that arterial ECM is subject to cyclic, time-dependent loading, we sought to characterize the viscoelastic, time-dependent mechanical properties of *COL3A1*^*+/G939D*^ CDM. We indented each sample to an indentation depth of 300 nm at a loading rate of 3000 nm/s and held this displacement for 5 seconds with the same spatial pattern of indentation as above (**Fig. S2**). Resulting load vs. time curves were analyzed to determine peak load, steady-state load, and half-time of relaxation (**Fig. 5e, Fig. S8**). We found that *COL3A1*^*+/G939D*^ CDM was characterized by a lower magnitude steady state load to peak load ratio (**Fig. 5f**) and that the half time of relaxation was significantly longer than healthy age/sex matched control (**Fig. 5g**). This stress relaxation behavior was specific to *COL3A1*^*+/G939D*^ CDM, as CDM generated from other vEDS genotypes were characterized by parameters similar in magnitude to respective controls (**Fig. 5f**,**g**). Interestingly, the trends in stress relaxation behavior correlated with CDM glycosaminoglycan (GAG) content, with *COL3A1*^*+/G939D*^ CDM characterized by an increased concentration of GAG compared to other *COL3A1*^*+/mutant*^ CDM and neonatal healthy control (**Fig S7e**).

### 3.7 Endothelial cell migration on CDM

To determine the functional and pathological consequences of the altered mechanics of *COL3A1*^*+/G939D*^ CDM, we plated HAECs onto healthy CDM and *COL3A1*^*+/G939D*^ CDM and monitored migration over 7 hours by imaging a fluorescently conjugated VE-cadherin antibody added to the live cells (**Fig 6a**). Through visualization of the migration tracks of these cells, we found that HAECs did not migrate as far from their starting point on *COL3A1*^*+/G939D*^ CDM as compared to healthy CDM (**Fig 6b**). We found that HAECs on *COL3A1*^*+/G939D*^ CDM also displayed lower migration speeds compared to those on healthy CDM (**Fig 6c**). In concordance with these findings, the mean straight-line speed and persistence were lower on the *COL3A1*^*+/G939D*^ CDM compared to the healthy CDM (**Fig 6d**,**e**). Overall, these results indicate that endothelial cells are sensitive to differences in the biochemical and biophysical properties of CDM generated from healthy fibroblasts as compared to CDM generated from the *COL3A1*^*+/G939D*^ donor, which is characterized by a decreased Young’s modulus and undergoes more significant load relaxation than CDM from healthy donors (**Fig. 5**).

**Fig. 6.**
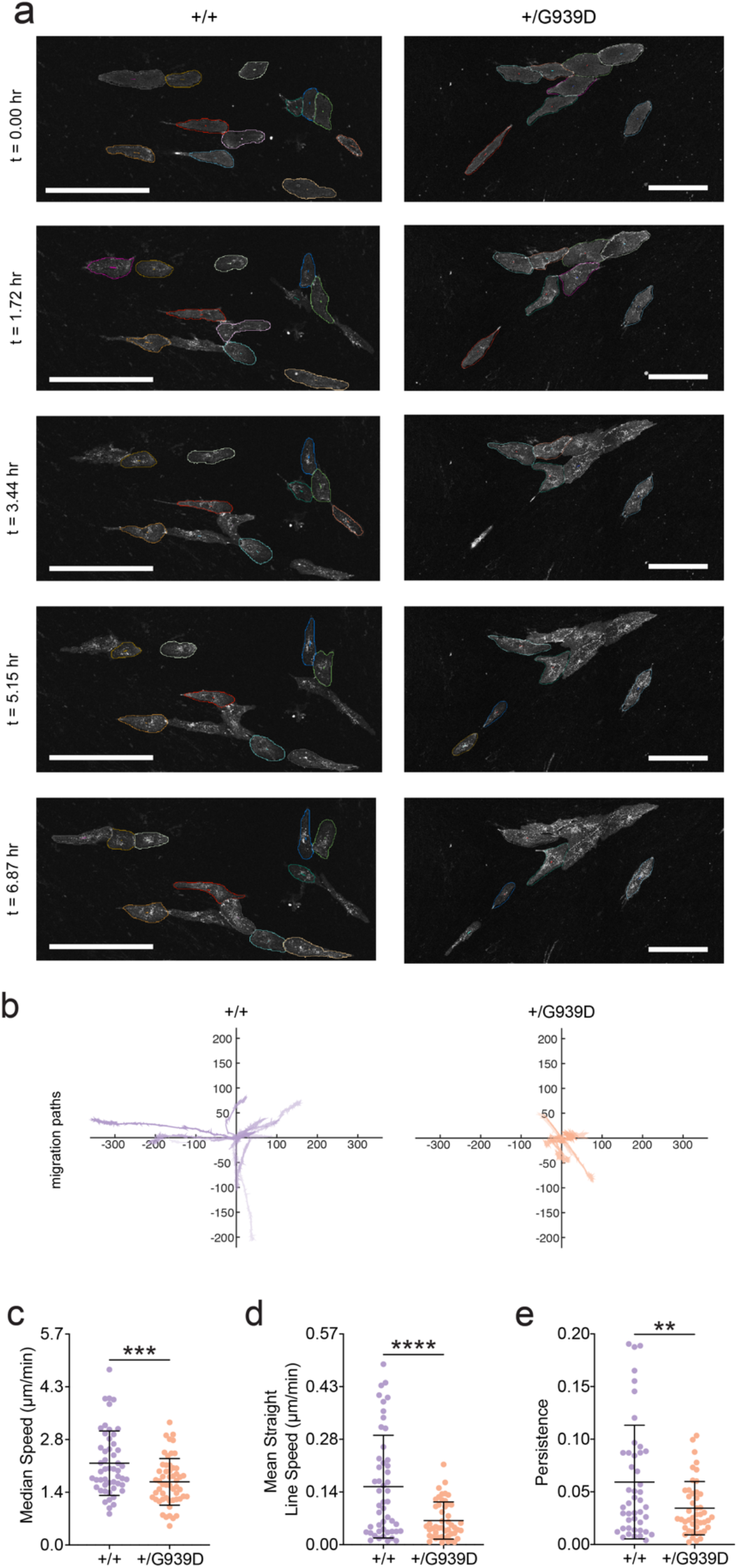
Endothelial cell migration is differentially regulated on +/G939D CDM. **a** Representative images of HAECs cultured on COL3A1^+/+^ CDM and COL3A1^+/G939D^ CDM throughout migration time course. HAECs labeled with AlexaFluor-647 anti-human VE-Cadherin antibody. Cells are shown overlayed with outline used in migration tracking. Scale bar = 200 µm. **b** Migration tracks for individual cells shown as (x,y) coordinates from origin point of migration. Migration characteristics of HAECs cultured on +/+ and +/G939D demonstrate a reduction of (**c**) median migration speed, (**d**) the mean straight line migration speed, and (**e**) the persistence. Statistics shown are a one-way ANOVA with a Tukey post-hoc test (*P<0.05, **P<0.01, ***P<0.001, ****P<0.0001), all plots are mean ± standard deviation.

## 4. Discussion

The severity of vEDS in terms of clinical complications and age of first clinical event varies with the type of *COL3A1* variant, and patients with variants resulting in glycine substitution present with more severe disease characteristics than the other variants examined here [44, 45]. However, it remains unclear how these variants contribute to varied clinical presentation and outcomes. The approach advanced in this study is the first to generate matrix directly from primary vEDS patient cells, which not only allows full proteomic characterization of how vEDS ECM differs in composition from healthy, but also allows functional studies to relate changes in composition to mechanical properties that have long thought to drive the clinical phenotypes. Using ECM produced by primary donor fibroblasts, we demonstrate that variants in *COL3A1* impact the production of type III collagen specifically (**Fig 1**), and the structure (**Fig 2**), composition (**Fig 3**), and mechanical properties (**Fig 5**) of the ECM more broadly. In particular, this is the first investigation of time-dependent mechanical properties in vEDS, and the findings suggest that ECM mechanical time-dependence may play a role in disease severity. Furthermore, in nearly every analysis conducted in this study, there was variation among the donors that was dependent upon their genotype when compared to healthy controls, suggesting that effective intervention to prevent disease progression could depend upon and should be tailored to the patient genotype.

A prevailing hypothesis in the pathogenesis of vEDS is that collagen III acts principally through covalent interactions with other collagens, including collagen I and collagen II [17, 22, 48]. Recent evidence in support of this hypothesis includes differential changes in collagen fibril diameter in healthy and heterozygous mice, which are characterized by a wider distribution of collagen fibril diameters, trending toward larger fibrils in cartilage [17] and skin [49]. In contrast to the observations in these murine models, we find that patient donor cells generate ECM with smaller fibrils and lower pore sizes when compared to healthy ECM (**Fig. 2**). While the absolute difference in fiber alignment and anisotropy of cell-derived ECM and that seen in connective tissue is likely attributable to tissue-scale assembly and remodeling mechanisms lacking in our reductionist model, it is also worth noting that in the haploinsufficient mouse model used to study COL3A1 in cartilage, a 50% reduction in *COL3A1* transcript expression was observed compared to healthy [23]. This is markedly different from the primary patient fibroblasts, which express *COL3A1* at similar or greater levels than healthy (**Fig. 1**), consistent with previous studies using donor fibroblasts [16]. Thus, our results broadly support that the causal mutations used in this study impact ECM assembly but further suggest the possibility for alternative mechanisms that regulate matrix structure than the previously suggested dose-dependent effects of *COL3A1* on collagen I assembly [21]. Furthermore, the glycine substitution variant demonstrated a similar rate of overall ECM production and collagen I content compared to the healthy control (**Fig. 2**), in contrast to murine models of *COL3A1* glycine substitutions where collagen I content in the adventitial wall was reduced [49, 50]. These results suggest that the effects of *COL3A1* mutations on matrix synthesis could differ in mouse and humans, as supported by pathway analysis demonstrating differences in the molecular pathways of key genes that regulate *COL3A1* expression in mouse and humans [17].

To determine biochemical changes in the deposited ECM that accompany changes in structure, we developed a pipeline for proteomic characterization of constituent ECM proteins deposited by primary donor fibroblasts (**Fig. 3**). We observed a decrease in abundance of many collagen chains, including COL1A1 and COL1A2, in CDM from *COL3A1*^*+/IVS40-1G>A*^ and *COL3A1*^*+/766delA*^ donors (**Fig 3**), consistent with picrosirius red staining (**Fig. 2**). Fibulin-1 (FBLN1) was found to be downregulated in all patient CDM samples (**Fig 3**), which is notable due to previous observations that lower expression of fibulin-1 has been seen to increase aortic stiffness in humans [51]. Fibulin-4 (EFEMP2), which was also downregulated in all patient CDM samples (**Fig 3**), is known to impact arterial mechanical properties by interacting with LOX to crosslink collagen and elastin [52], and loss of fibulin-4 has been shown to alter collagen fibril size [53] and to impact the viscoelastic properties of the aorta [52]. While there was no consistent trend among all LOX proteins seen in our proteomic data set (Fig 3), LOX and LOX4 were consistently downregulated among all patient samples. Together these results suggest that the effects of *COL3A1* mutations on aneurysm susceptibility could be driven by altered expression of fibulin and LOX and resulting impacts on arterial elasticity.

Interestingly, tenascin-X was upregulated in all *COL3A1*^*+/mutant*^ CDM compared to healthy CDM (**Fig. 3**). Tenascin-X is a glycoprotein with elastic properties [54] that binds to collagens and other ECM proteins and has been shown to increase the elastic modulus of collagen I hydrogels under compression in a dose-dependent manner [55]. Deficiency in tenascin-X can induce a recessive form of classical EDS [56], and tenascin-X has also been shown to activate latent TGF-β1 and regulate epithelial-to-mesenchymal transition (EMT) in epithelial cells [57]. Furthermore, circulating TGF-β1 has been shown to be upregulated in patients with vEDS [58], and while the role of TGF-β1 in aneurysm progression is complex with some models demonstrating a role in aneurysm progression and other demonstrating a role in healing [59], previous studies demonstrate the activity of TGF-β1 in driving EMT is dependent on the stiffness of the underlying matrix [60]. Thus, the concomitant changes in matrix stiffness (**Fig. 5**) and biochemical composition could play a role in vEDS pathogenesis and aneurysm progression and represent an opportunity for further exploration.

To determine the functional consequences of these changes in matrix constitution and structure, we measured the time required for embedded fibroblasts to contract cell-generated matrix. Interestingly, we found that the *COL3A1*^*+/IVS40-1G>A*^ and *COL3A1*^*+/766delA*^ donor cells took significantly longer to contract the cell-generated matrix, despite similar levels of traction force generation by cells seeded in 2D and 3D, in contrast to cells from the donor with the glycine-substitution mutation, which contracted the matrix at similar rates to healthy control (**Fig. 4**). Notably, the rate of contraction correlated with collagen content and fibril diameter, with lower collagen content and smaller fibril sizes (**Fig. 2**) corresponding to slower rates of contraction, consistent with previous observations that collagen content is a key driver of fibroblast-mediated tissue contraction [61]. To determine whether these changes in matrix structure and composition directly impacted mechanics of the ECM, we conducted indentation mechanical analysis on CDM. It is worth noting that we observed significant increases in ECM modulus with donor age and as a function of donor biological sex for both healthy and vEDS donors (**Fig. S6**). Furthermore, the ECM generated from donor fibroblasts with glycine substitution mutation demonstrated a significantly decreased Young’s modulus (**Fig. 5**). While these are the first direct measurements of the mechanical properties of isolated vEDS ECM, the tensile force at rupture has been measured previously for murine aortic explant rings. Aortic segments from heterozygous loss of function mice (*COL3A1*^*m1Lsmi/+*^) were characterized by a reduced maximum tensile force compared to healthy control [62], which was consistent with our overall findings for *COL3A1*^*+/mutant*^ ECM.

We next sought to characterize the viscoelastic mechanical properties of the ECM through a load relaxation study, and while the CDM from *COL3A1*^*+/IVS40-1G>A*^ and *COL3A1*^*+/766delA*^ donors demonstrated similar properties to age- and sex-matched healthy controls (**Fig. 5**), the CDM from *COL3A1*^*+/G939D*^ donor fibroblasts was characterized by decreased steady state/peak load ratio and an increased time constant of load relaxation (**Fig. 5**). Interestingly, CDM from this donor was also characterized by increased proteoglycan content (**Fig. 3**) and total GAG content (**Fig. S7e**), and consistent with these results, GAG content has been shown to contribute to time-dependent mechanical properties through modulation of ECM permeability [63]. GAG levels have also been shown to affect the mechanical properties of arterial vessel walls through interactions in collagen fibers [64]. In addition to total GAG content, proteomic characterization revealed a higher abundance of proteoglycans (**Fig 3**) in the CDM from the fibroblasts with the glycine substitution mutation and specific proteoglycans such as biglycan (BCAN) are known to impact stress relaxation [65]. Recent work has demonstrated that the viscoelastic properties of the ECM govern cell migration and that when cultured on materials with equivalent elastic moduli, cells migrated more slowly and extended fewer protrusions on ECM with longer time-constants for stress relaxation [66]. Consistent with these results, we observed that HAECs migrated more slowly with lower persistence when cultured on ECM from *COL3A1*^*+/G939D*^ fibroblasts (**Fig. 6**). Given reported correlations between reestablishment of the endothelial lining and the stabilization of aneurysms [67], these results suggest that the viscoelastic mechanical properties of the ECM could act as an inhibitor of aneurysm stabilization through effects on endothelial cell migration.

An overarching challenge to the treatment of vEDS is lack of clarity regarding how mutations in *COL3A1* render the arterial vasculature susceptible to aneurysm formation. The prevailing hypothesis has been that the dominant negative effect of *COL3A1* variants on collagen III assembly results in compromised mechanical properties of the arterial wall that renders vessels susceptible to rupture. Here, we interrogate this hypothesis using a novel method for studying ECM generated by patient donor cells, and we demonstrate that the effects of causal mutations on ECM biochemical and biophysical properties depend on genotype, and that the underlying mechanism that results in aneurysm formation and progression could depend on the specific type of causal mutation. Furthermore, our results suggest that matrix constituents other than type III and type I collagen could provide targets for therapeutic intervention for vEDS and other diseases in which *COL3A1* variants or expression levels play a role in pathogenesis, including pulmonary arterial hypertension [68], pulmonary fibrosis [69], and various solid tumor types [70, 71]. The observation that EC migration is regulated by *COL3A1* mutations highlights a potential role of endothelial cells in vEDS disease progression and motivates further investigation into the role of the vascular endothelium in disease progression.

## Acknowledgements

This work was supported by the National Institutes of Health (R35GM142944) and by the American Heart Association (CDA857738). E.L.D. acknowledges financial support of the National Institutes of Health through the Integrative Vascular Biology Training Program (T32HL69768) and a Ruth L. Kirchstein predoctoral individual fellowship (F31HL162462). W.Y.A. is supported by grants from the CLOVES Syndrome Community and the Eshelman Institute for Innovation. The mechanical testing was performed in the Chapel Hill Analytical and Nanofabrication Laboratory, CHANL, a member of the North Carolina Research Triangle Nanotechnology Network, RTNN, which is supported by the National Science Foundation (ECCS-2025064), as part of the National Nanotechnology Coordinated Infrastructure, NNCI. The proteomics work was performed in part by the Molecular Education, Technology and Research Innovation Center (METRIC) at NC State University, which is supported by the State of North Carolina. We thank Dr. Matthew Kutys (Department of Cell and Tissue Biology, University of California, San Francisco) for helpful discussion regarding preparation of cell derived matrix.

