## Supplementary Information for "Patient-derived extracellular matrix demonstrates role of COL3A1 in blood vessel mechanics"

*SUPPLEMENTAL METHODS*

*S.1 Fibroblasts*

Specific information regarding the primary human fibroblasts used in this study are found in Table S1.

Table S1. Cell source, donor characteristics, and genotype for fibroblasts.

| Abbreviation | Age | Sex | Mutation in COL3A1 | Source | Catalogue Name |
|---|---|---|---|---|---|
| COL3A1$^{+/IVS40\text{-}1G>A}$ | 41 yr | M | IVS40-1G>A (PTC) | Corriel Inst. | GM21788 |
| COL3A1$^{+/766delA}$ | 47 yr | F | 766delA | Corriel Inst. | GM22609 |
| COL3A1$^{+/G939D}$ | 36 yr | F | Gly939ASP | Corriel Inst. | GM22779 |
| WT (WS 1) | Gest. | F | Not Seq. | ATCC | WS 1 |
| HDF 1650 | 37 yr | F | Not Seq. | Corriel Inst. | GM01650 |
| HDF 1653 | 37 yr | M | Not Seq. | Corriel Inst. | GM01653 |
| HDF 275 | 42 yr | M | Not Seq. | Corriel Inst. | GM00275 |
| HDF 5879 | 48 yr | F | Not Seq. | Corriel Inst. | GM05879 |

*S.2 qRT-PCR*

For qRT-PCR, a custom two-step hold stage defined by 50 °C for 2 min followed by 95 °C for 10 min, a two-step PCR stage of 40x cycles of 95 °C for 15 s followed by 60 °C for 1 minute, and a three step melt curve stage of 95C for 15 s, followed by 60 °C for 1 minute, followed by a 0.05 C/s ramp up to 95C, which is then held for 15 s. All ramps were set to 1.6C/s to move between temperatures unless otherwise specified. Specific gene targets were detected with the primer sequences listed in Table S2. Gene expression was calculated using -delta(delta(CT)) method where dCT equals the difference in CT value between COL3A1 gene and a housekeeping EIF4A2 gene.

Table S2. Primer sequences for gene expression analysis.

| Primers used for qRT-PCR | |
|---|---|
| **Gene** | **Sequence** |
| COL3A1_F | 5’-TCTGCCATCCTGAACTCAAGA-3’ |
| COL3A1_R | 5’-TGCATGTTTCCCCAGTTTCC-3’ |
| EIF4A2_HSA_F | 5’-CGGGATTGATGTGCAACAAGTG-3’ |
| EIF4A2_HSA_R | 5’-ATGGGCATCTCCTCCACTGTAG-3’ |

*S.3: Western Blot*

Thawed lysate was run using NuPAGE 4-12% Bis-Tris gels (Invitrogen, NP0323BOX) at 150V for 1 hour in NuPAGE MOPS Running Buffer (Invitrogen, NP0001) and then transferred onto PVDF membrane (Invitrogen, IB24001) using an iBlot system (Invitrogen). Blots were then blocked with 5% milk in TBST for 1 hour, washed 3x with TBST, and incubated with primary antibodies (Table S3) in 5% BSA in TBST overnight at 4 ˚C. The next day, blots were washed three times with TBST and incubated with secondary antibodies in 5% milk in TBST for one hour at room temperature. Blots were then washed three times with TBST and once with DI-$H_20$. The blot was cut in half to divide COL3A1 and GAPDH bands. Finally, the bands were developed using Femto ECL kit (ThermoFisher, 34094) for 5 minutes (COL3A1) or 5 s (GAPDH) and imaged with an iBright 1500 imaging system (Invitrogen). Blots were quantified using the raw chemiluminescent images in FIJI. Band intensities for both COL3A1 and GAPDH were then normalized to the first row of the gel and then COL3A1 intensity was divided by GAPDH intensity for each lane.

Table S3. Antibody information for Western Blot.

| Protein | Primary Antibody | | Secondary Antibody | |
|---|---|---|---|---|
| | Supplier Cat # | Dilution | Supplier Cat # | Dilution |
| COL3A1 | Santa Cruz SC-271249 | 1:200 | Goat-anti-Mouse Invitrogen 31466 | 1:2500 |
| GAPDH | Cell Signaling Tech 2118S | 1:1000 | Goat-anti-Rabbit Invitrogen 21040 | 1:5000 |

*S.4 TFM image analysis*

Traction force microscopy images were analyzed using the traction force microscopy package published by the Danuser lab [1], as described previously [2]. Briefly, images were processed into single frames and passed through the MATLAB GUI. Efficient subpixel registration was used for drift correction. Displacement field calculations assumed no outward deformations and high-resolution subsampling of beads. Bead correlation used a template size of 21 pixels with a maximum displacement of 20 pixels. Fourier transform traction cytometry (FTTC) was used with a constant regularization factor (0.0001), Young's modulus of 8 kPa, and gel thickness of 135 $\mu$m to reconstruct traction forces from the computed displacement fields. Custom MATLAB scripts were used to calculate strain energy, strain energy density, average traction, and maximum traction from each traction field as defined in Butler et al., 2002 [3]. Cell spread area was computed in FIJI with manual cell segmentation of the first image frame.

*S.5 Proteomics pipeline*

CDM samples were generated in Mattek glass bottom dishes or 6 well plates as described above. Samples were then sent to METRIC Core Facility at NCSU for processing and liquid chromatography-mass spectrometry (LC-MS) analysis. Briefly, samples were washed twice with DI-$H_2O$ and scraped from each plate into a microcentrifuge tube with 500 µL DI-$H_2O$. Samples were then homogenized with a Genogrinder for eight 30 s agitation cycles at 1400 rpm with a 30 s rest period between agitations using 2.8 mm ceramic beads. A 200 µL aliquot of each sample was treated with DTT and IAA to alkylate cysteines, then digested with tryspin/lys-C. Peptides were separated from undigested material using 10 kD MWCO filters. Undigested collagen material that remained on the filter was further treated with elastase for 4 hrs at 37 $^{o}$C. Peptides were separated from digestion enzymes by centrifugation. Recovered peptides from both enzyme treatments were separately analyzed by LC-MS/MS.

To perform LC-MS/MS, each sample was injected on an Easy-Nano-1200 nanoLC system (Thermo Scientific, San Jose, CA, USA) interfaced with an Orbitrap Exploris 480 (Thermo Scientific) Mass Spectrometer. Chromatography employed a 0.075 mm × 20 mm $C_{18}$ trap column with particle size of 3 µm (Thermo Scientific Accclaim PepMap$^{TM}$ 100, Part # 164946) in line with a 0.075 mm × 250 mm $C_{18}$ nanoLC analytical column with particle size of 2 µm (Thermo Scientific PepMap$^{TM}$, Part # ES902). A solvent gradient of water containing 2% acetonitrile, 0.1% formic acid (MPA) and acetonitrile containing 20% water, 0.1% formic acid (MPB) was used. MPB was held at 5% for 2 min, increased to 25% over 75 min, increased to 40% over 10 min, increased to 95% in 1 min, and was held at 95% for 17 min. Mass spectrometer parameters are listed here: 1.8 kV positive ion mode spray voltage, ion transfer tube temperature of 275 °C, master scan cycle time of 2.5 s, m/z scan range of 300 to 1,500 at 120 K resolution, 300% normalized AGC Target, 120 ms maximum $MS^1$ injection time, RF lens of 40%, 15 K mass resolving power for data-dependent $MS^2$ scans, 1.5 m/z isolation window, 30% normalized HCD collision energy, 100% normalized AGC Target, 21 ms maximum injection time and dynamic exclusion applied for 20 s periods.

*S.6 Proteomics data analysis*

Mass spectrometry data were processed with Proteome Discoverer 2.5 software (PD, Thermo Scientific, San Jose, CA) using a Homo sapiens protein database (Taxon 9606) obtained from Swiss-Prot (42,253 sequences). Two raw files corresponding to injections of separate enzyme digestions for each sample were combined as fractions and processed as a single sample by PD. A custom cleavage reagent was made that combined elastase and trypsin cleavage sites at alanine, isoleucine, lysine, leucine, arginine, and valine at the c-terminus with proline as an inhibitor. Searches included a custom contaminants database to identify human keratin and reagent enzyme peptides. Two SEQUEST HT nodes were put in linear sequence. The first node used the custom cleavage enzyme (full) and the second used the default trypsin enzyme (full). All other parameters were the same for both search nodes. SEQUEST HT was set up as follows: maximum of 3 missed cleavage sites; minimum peptide length of 6 amino acids; 5 ppm precursor mass tolerance; 0.02 Da fragment mass tolerance; maximum of 8 equal dynamic modifications, which were oxidation of lysine, methionine, and proline; static carbamidomethylation of cysteine. After the first search node, peptides were validated by Percolator with q-value set to 0.05 and strict false discovery rate (FDR) set to 0.01. All peptides with a confidence level "worse than high" were allowed to pass to the next SEQUEST node. The second Percolator node q-value and FDR were 0.05 and 0.01, respectively.

Processed data was analyzed using label-free quantification in Proteome Discoverer to determine the UniProt IDs of significantly up- and downregulation of proteins in COL3A1$^{+/-}$ samples in relation to COL3A1$^{+/+}$ (WS1) samples. Lists of up- and downregulated proteins UniProt IDs were then individually input into PANTHER GO TERMS analysis for molecular function using the Homo Sapiens Reference Database. Significant molecular functions were extracted and plotted as function of FDR.

*S.7 Modified Hertzian model for elastic mechanics*

First, raw load vs. indentation data were uploaded into MATLAB using a custom written script. Then, load vs. indentation curves were visually inspected to remove curves resulting from indentation of glass or incomplete data acquisition (**Fig. S1**). Then, curves were sectioned from the initial indentation depth to the maximum indentation depth to isolate the loading curve, and the contact point was manually selected for each curve by determining the point where a change in concavity of the curve occurs. Next, the curves are fit using a nonlinear least-squares fitting method to a modified Hertzian model that accounts for the finite thicknesses of the CDM (Eqs. S1-2) [4]:

$$F = \frac{16E}{9} R^{1/2}\delta^{1/2}[1 + 1.133\chi + 1.283\chi^2 + 0.769\chi^3 + 0.0975\chi^4] \quad (S1)$$

$$\chi = \sqrt{R\delta}/h \quad (S2)$$

Where $\mathrm{R}$ = cantilever tip radius, $\mathrm{F}$ = load, $\delta$ = indentation depth, and $\mathrm{h}$ = sample height. The sample height was determined by the average CDM thickness for each genotype as determined by reflectance microscopy on day seven.

*S.8 Determining fit parameters for modified Hertzian model*

Raw data was imported into MATLAB trimmed to include only the loading curve (**Fig. S8a**). The modified Hertzian model equation (Eqs. S1-2), was fit to the loading curve over a range of fit windows for each curve. Each fit window, *n*, spanned from the contact point, defined as displacement of 0, to displacement of 0 + 100 * *n,* with $n_{max}$ defined as the window that spanned from 0 to the displacement resulting in the peak load (**Fig. S8b**). In addition, different smoothing span increments were used to smooth the data using the 'sgolay' algorithm to remove noise (**Fig. S8b**). These analyses resulted in a Young's modulus value and $R^2$ value for each fit distance and smoothing span (**Fig. S8c**). A sensitivity analysis was then run to determine which smoothing factor and distance from the contact point give the best fit (**Fig. S8d**). The top half of $R^2$ values (greater than $R^2$ = 0.9 or lowering the threshold until at least 10 values were selected) were identified, the most prominent distance from the contact point among these values was selected, and the resulting Young's modulus values were ranked in order of magnitude from least to greatest. Finally, the percent difference between the Young's modulus values was assessed and if the difference in value is less than 5% then the Young's modulus value is selected as the best fit.

*S.2: Stress-Relaxation Data Analysis*

The raw data from stress-relaxation experiments were imported into MATLAB using a custom written script, and the characteristic decay curve was isolated by finding the peak load and extracting load vs. time data from peak load to 1 second after the peak (**Fig. S9a-b**). Curves that reflected indentation of glass or incomplete processing were discarded. Each curve was then normalized by dividing by the peak load value and smoothed to remove noise (**Fig S9c-d**). The steady-state load to peak load ratio was found by averaging the last 200 data points of the load vs. time curve to determine the steady state load (**Fig S9e**). The half-time was found by determining time at which the load reached a magnitude of 0.5 * (peak load – steady state load) (**Fig S8f**). This analysis pipeline was applied to each of the 25 curves for each sample and then averaged to obtain a characteristic curve for each sample (**Fig S9g-h**), and each sample curve was averaged to achieve characteristic values for each condition.

Table S5. Statistical results of Robust Linear Mixed Models

| ROBUST LMM ALL DATA | | | | | |
|---|---|---|---|---|---|
| | Estimate | Std Error | t value | p value | sig. |
| intercept | 1.07627 | 0.93878 | 1.146 | 2.61E-01 | ns |
| age | 0.10873 | 0.02189 | 4.967 | 3.03E-05 | **** |
| sex | 0.58696 | 0.21081 | 2.784 | 9.51E-03 | ** |
| vEDS | -0.55111 | 0.20648 | -2.669 | 1.25E-02 | * |

| **ROBUST LMMs Removing +/G393D and age-matched +/+ Data**<br>*Age v vEDS* | | | | | |
|---|---|---|---|---|---|
| | Estimate | Std Error | t value | p value | sig. |
| intercept | 5.53434 | 1.802034 | 3.071 | 0.00697071 | ** |
| age | 0.003726 | 0.039741 | 0.094 | 0.92640258 | ns |
| vEDS | 0.029766 | 0.259584 | 0.115 | 0.91005685 | ns |

| *Sex v vEDS* | | | | | |
|---|---|---|---|---|---|
| | Estimate | Std Error | t value | p value | sig. |
| intercept | 5.7132 | 0.24677 | 23.152 | 3.46E-14 | **** |
| sex | -0.02236 | 0.23845 | -0.094 | 9.26E-01 | ns |
| vEDS | 0.02604 | 0.25903 | 0.101 | 9.21E-01 | ns |

1

*SUPPLEMENTAL FIGURES*

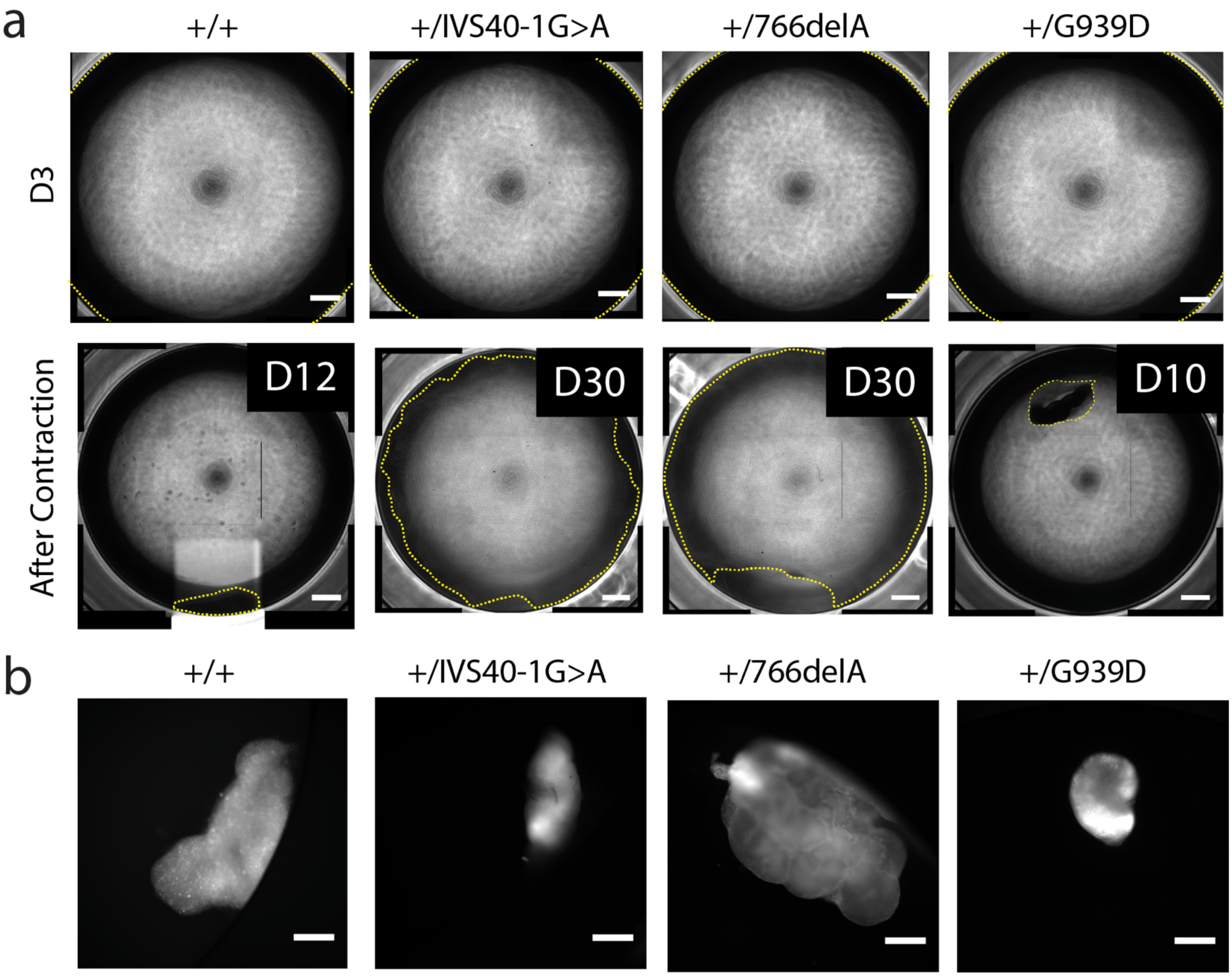


**Fig. S1** – **Cell sheet contraction. a** Cell sheet contraction visualized with phase contrast imaging on day 3 after plating (top) and the day of contraction (shown in top right corner). Yellow dotted line shows the edge of the cell sheet (scale bar = 1000 μm). **b** Micrographs of cell-laden ECM after contraction. Cells were stained with CellMask Deep Red (scale bar = 500 μm).

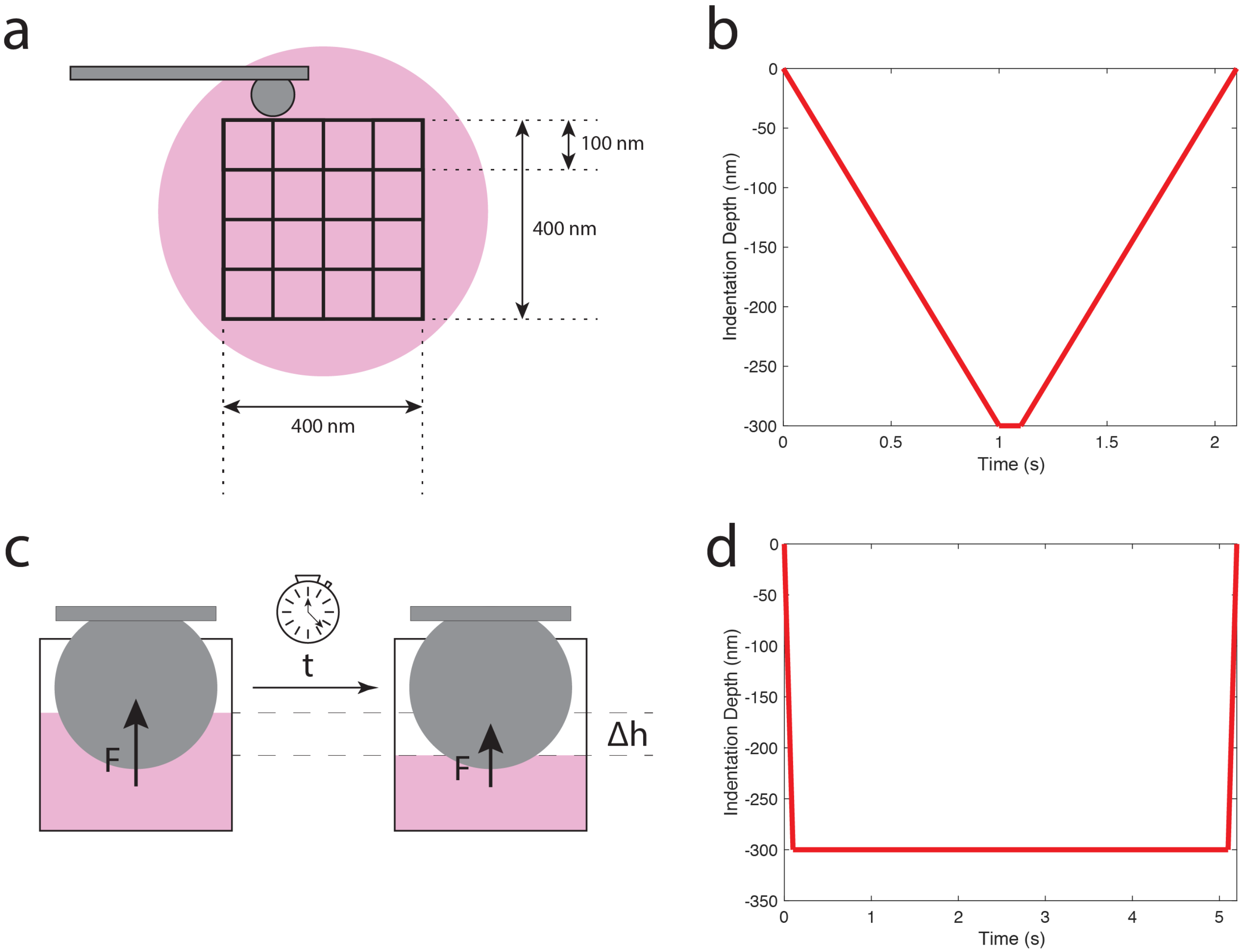


**Fig. S2** – **Nanoindentation Parameters**. (**a**) Map of indentation grid with distance labels shows the 25 positions collected during nanoindentation acquisitions for Young's modulus and load-relaxation measurements. (**b**) Indentation profile for Young's modulus nanoindentation acquisition. (**c**) Schematic of load-relaxation study where a constant indentation depth is maintained over time and load is recorded as a function of time. (**d**) Indentation profile for stress-relaxation nanoindentation acquisition.

*Supplemental Data*

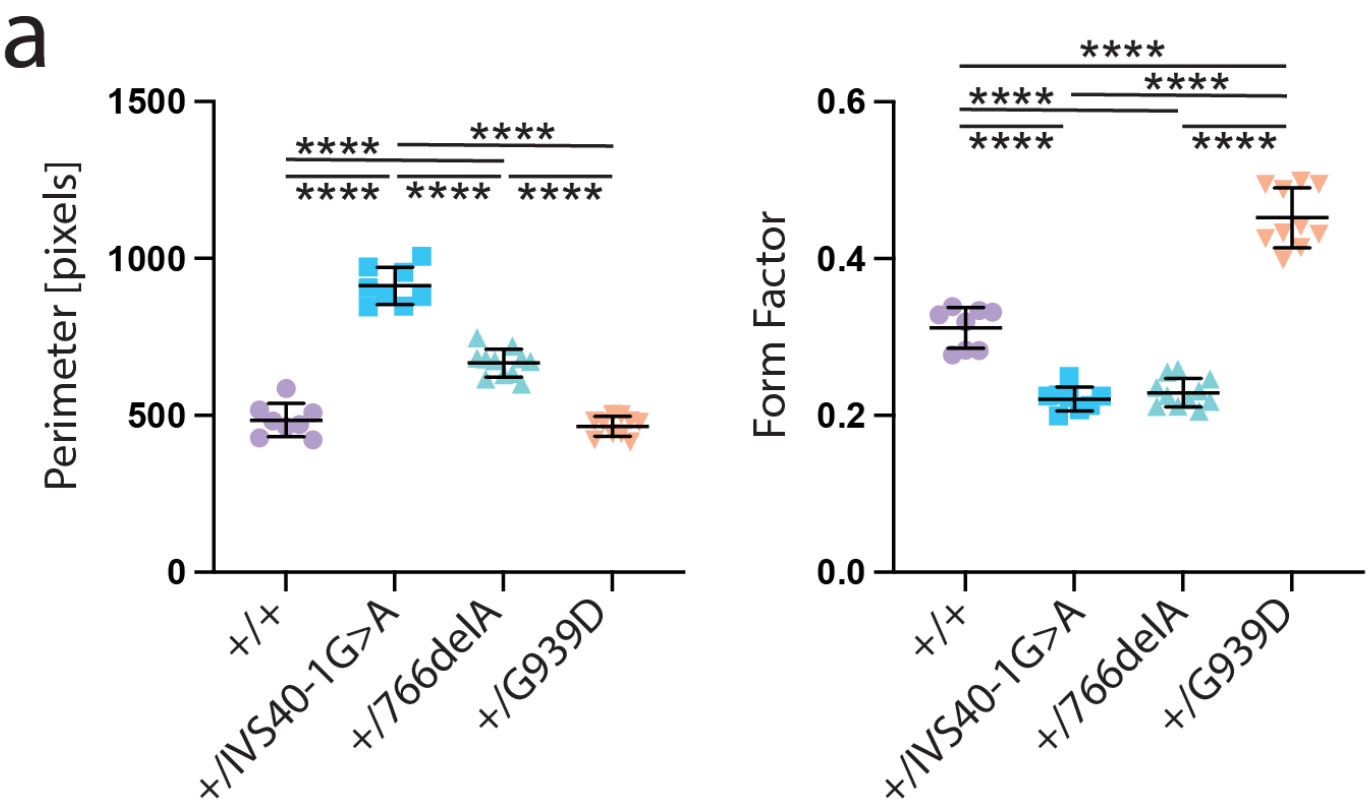


**Fig. S3 – Cell shape characteristics for vEDS donor and wild-type fibroblasts**. Perimeter and form factor of cells, quantified from phalloidin-stained fibroblasts. Each data point is the average perimeter or form factor of all cells in an image from an independently prepared sample (*n* > 8 independent samples, statistics shown are a one-way ANOVA with a Tukey post-hoc test. All plots, mean ± standard deviation, *P<0.05, **P<0.01, ***P<0.001, ****P<0.0001).

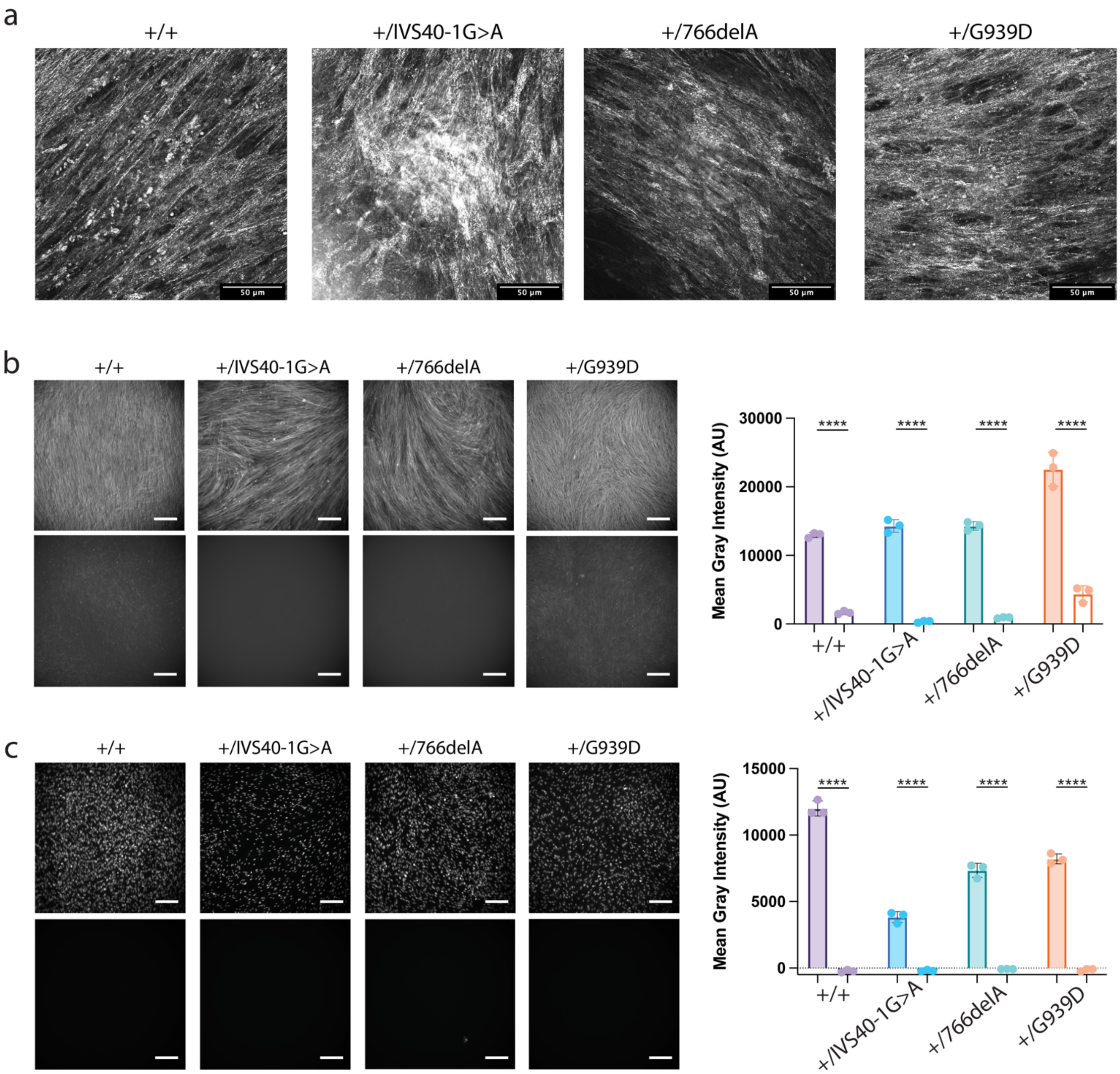


**Fig. S4 – CDM generated from patient donors was effectively decellularized**. (**A**) Confocal reflectance microscopy images captured at 60X with a 488 nm laser (scale bar = 50 μm). CDM stained with (**b**) phalloidin and (**c**) DAPI before (top) and after decellularization (bottom, scale bar = 200 μm). Mean gray intensity for n=3 biological replicates for each condition with (solid bars) and without (open bars) decellularization. Data is represented as mean +/- standard deviation and statistics shown are a two-way ANOVA with a Tukey post-hoc test, *P<0.05, **P<0.01, ***P<0.001, ****P<0.0001**.**

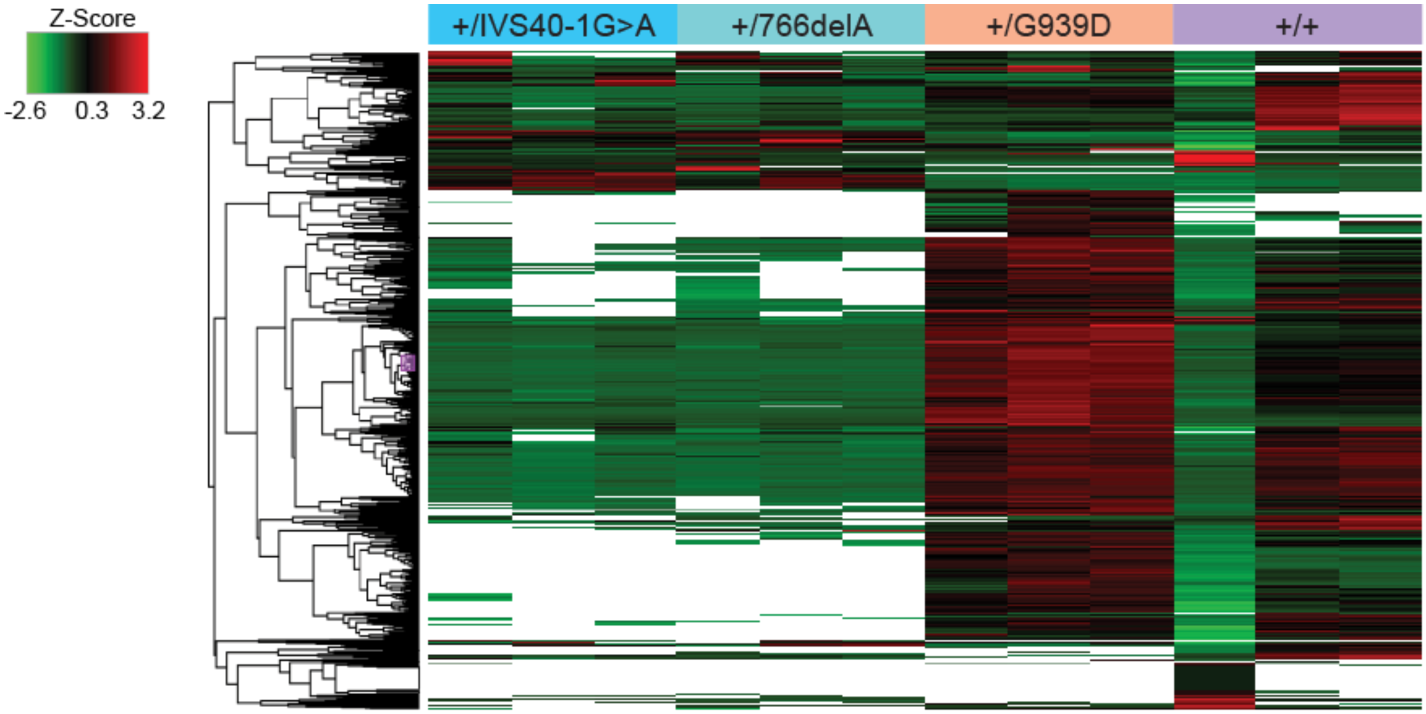


**Fig. S5 – Full Proteomics Heat Map.** Heat map of proteomic characterization of COL3A1$^{+/+}$ and COL3A1$^{+/mutant}$ CDMs. This map shows normalized abundance values for each replicate (n = 3 replicate per genotype) scaled to a z-score before hierarchical clustering using the Euclidian distance function and the complete linkage method (shown by the hierarchy tree on the left). White spaces indicate that the protein was not found in that sample.

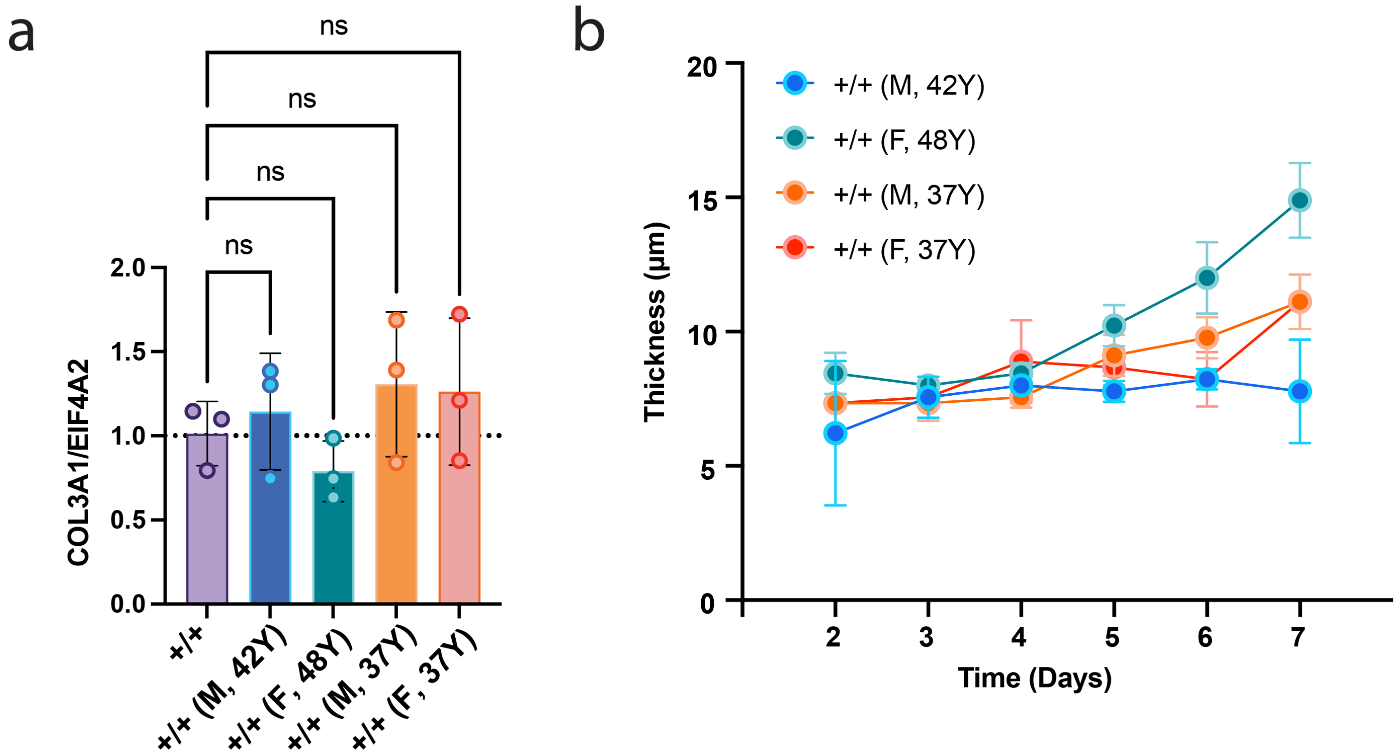


**Fig. S6 – Adult healthy HDF characterization.** (**a**) Relative gene expression of *COL3A1* to *EIF4A2* gene expression measured by qRT-PCR in cultured fibroblasts from healthy adult donors (*n* = 3 independent lysates, statistics shown are a one-way ANOVA with a Tukey post-hoc test). (**b**) Thickness of deposited ECM as measured daily by confocal reflectance microscopy at 488 nm (line indicates mean and corridor indicates standard deviation, as measured from *n* = 3).

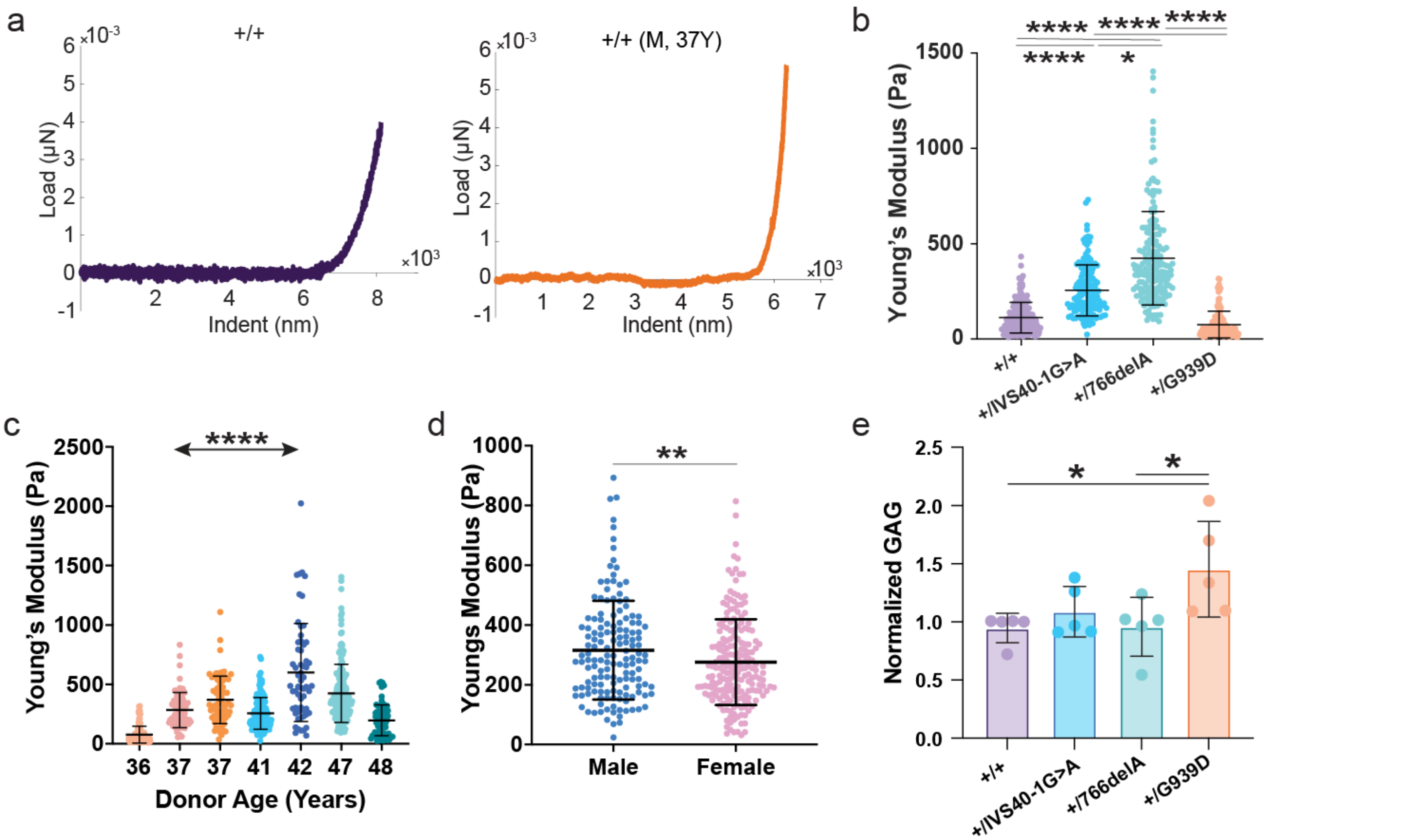


**Fig S7 – CDM elastic mechanical properties and glycosaminoglycan concentration.** (**a**) Representative load v. indentation curves for neonatal +/+ CDM and +/+ (M, 37Y) CDM (an age, but not sex matched control for +/G939D). (**b**) Young's modulus as computed from indentation curves for +/mutant CDM plotted against Young's modulus of +/+ neonatal CDM. (**c**) Young's modulus data for +/mutant and age/sex matched +/+ CDM, categorized by age. Data is presented as mean +/- standard deviation, each dot is an individual Young's modulus measurement. (**d**) Young's modulus data for +/mutant and age/sex matched +/+ CDM, categorized by sex. (**e**) Total GAG content in CDM normalized to the healthy control content (**c**) Young's modulus data for +/IVS40-1G>A and +/766delA and age/sex matched +/+ CDM. Data is presented as mean +/- standard deviation, each dot is an individual Young's modulus measurement. Statistics shown for panels b and e are a one-way ANOVA with Tukey post-hoc test (*P<0.05, **P<0.01, ***P<0.001, ****P<0.0001). Statistics shown for panels c and d are a robust linear mixed model assuming the same degrees of freedom as a traditional linear mixed model (*P<0.05, **P<0.01, ***P<0.001, ****P<0.0001).

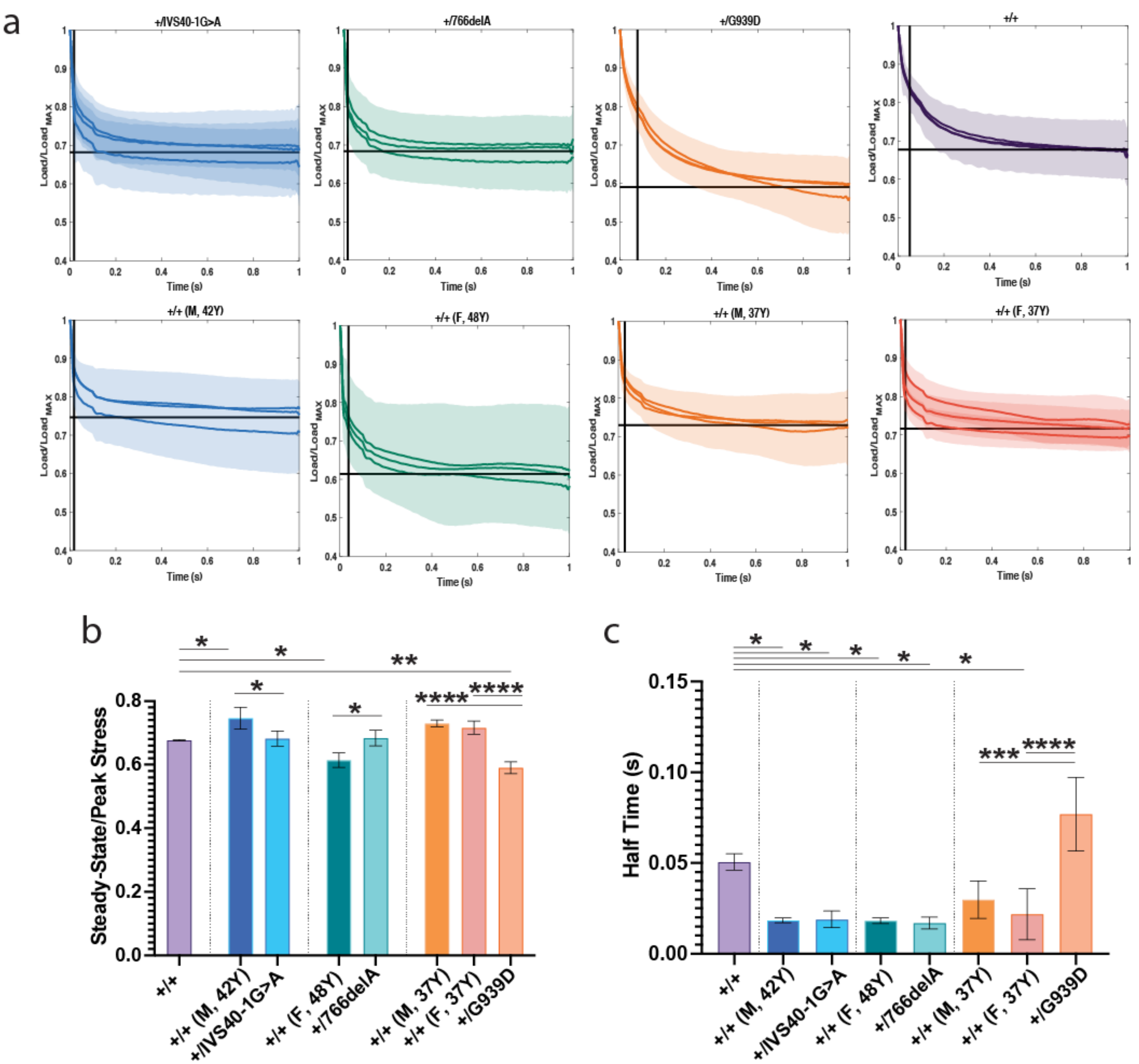


**Fig. S8 – Full Stress-Relaxation Data.** (**a**) Load v. time curves for all CDMs. Three samples (n = 3 for all conditions) are shown as average of 12-25 individual curves. Corridor shows standard deviation from mean (solid colored line) for each sample. Black horizontal lines indicate the average steady-state/peak stress for all samples. Black vertical lines indicate the average half time for all samples. (**b**) Steady-state/peak stress data plotted as mean +/- standard deviation. Statistics shown are a one-way ANOVA with a Tukey post-hoc test (*P<0.05, **P<0.01, ***P<0.001, ****P<0.0001). (**c**) Half time relaxation data plotted as mean +/- standard deviation. Statistics shown are a one-way ANOVA with a Tukey post-hoc test (*P<0.05, **P<0.01, ***P<0.001, ****P<0.0001).

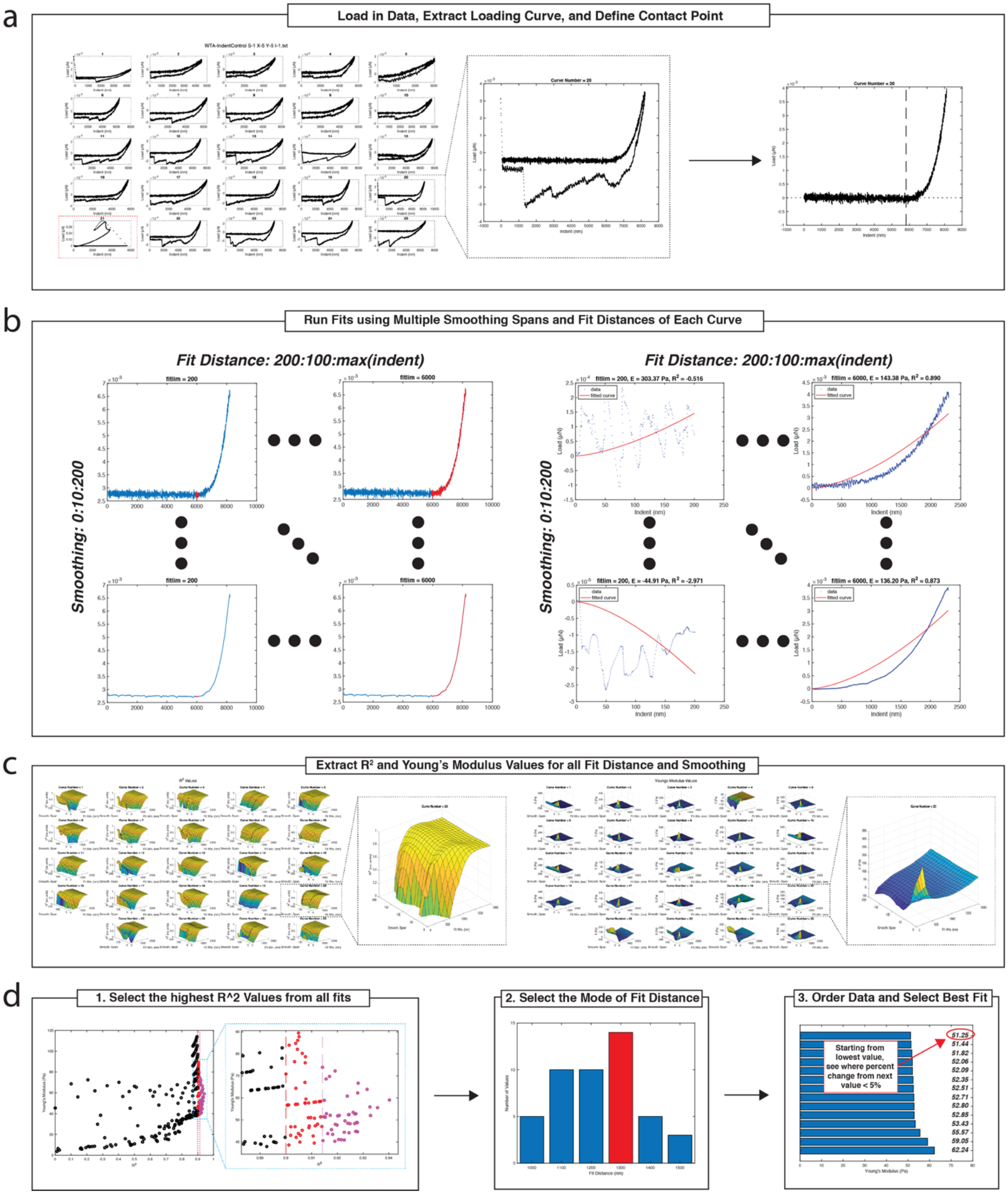
a
Load in Data, Extract Loading Curve, and Define Contact Point
Curve Number = 20
Indent (nm)
Load (μN)
b
Run Fits using Multiple Smoothing Spans and Fit Distances of Each Curve
Fit Distance: 200:100:max(indent)
Fit Distance: 200:100:max(indent)
Smoothing: 0:10:200
Smoothing: 0:10:200
fitlim = 200
fitlim = 6000
fitlim = 200, E = 303.37 Pa, R² = -0.516
fitlim = 6000, E = 143.38 Pa, R² = 0.890
fitlim = 200, E = -44.91 Pa, R² = -2.971
fitlim = 6000, E = 136.20 Pa, R² = 0.873
data
fitted curve
Indent (nm)
Load (μN)
c
Extract R² and Young's Modulus Values for all Fit Distance and Smoothing
d
1. Select the highest R^2 Values from all fits
2. Select the Mode of Fit Distance
3. Order Data and Select Best Fit
Young's Modulus (Pa)
Number of Values
Fit Distance (nm)
Starting from lowest value, see where percent change from next value < 5%
51.25
51.44
51.82
52.06
52.09
52.35
52.51
52.71
52.80
52.85
53.43
55.57
59.05
62.24

1

**Fig. S9 – Young's Modulus Methodology.** (**a**) 25 raw curves are imported into MATLAB, and each curve is trimmed, and the contract point is defined based on a change in concavity of the indentation curve. (**b**) Each curve is trimmed to a variety of distances from the contact point (left and top, fit distance shown in red), and smoothing by averaging over windows of increasing size is applied (left and bottom, largest amount of smoothing shown on the bottom). Fits are applied for each of these variable values (right). (**c**) Young's modulus and $R^2$ values were extract for each fit distance and smoothing span applied to the raw data. (**d**) The matrix of values shown in (c) were then filtered through a series of rules to select the best fit for the data and extract the Young's modulus for each curve.

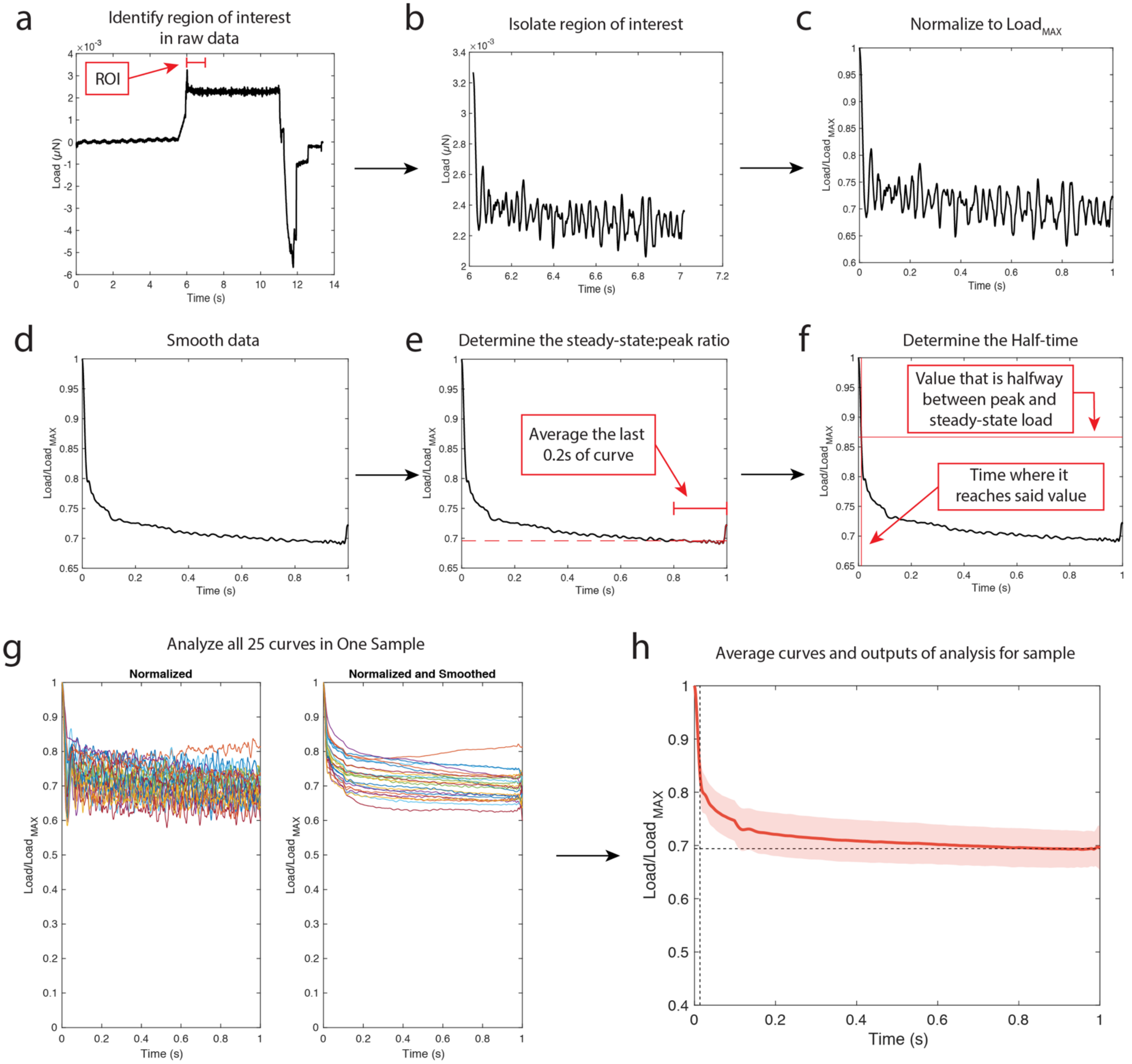


**Fig. S10 – Stress Relaxation Methods.** (**a**) Raw data was uploaded into MATLAB, and the region of interest was identified by finding the peak load value and (**b**) isolating the region spanning from peak load to 1 sec after peak load. (**c**) The trimmed curve was then normalized to the peak load value and (**d**) smoothed. The processed curves were used to extract the (**e**) steady-state/peak load ratio by averaging the last 0.2s of the curve and the (**f**) half-time by finding the time at which the decay in load has reach half of the steady state load. (**g**) 25 normalized and smoothed curves were (**h**) averaged for to achieve a mean load vs. time curve for each sample.
